# ERK-mediated mechanochemical waves direct collective cell polarization

**DOI:** 10.1101/2019.12.25.888552

**Authors:** Naoya Hino, Leone Rossetti, Ariadna Marín-Llauradó, Kazuhiro Aoki, Xavier Trepat, Michiyuki Matsuda, Tsuyoshi Hirashima

**Author notes:** Corresponding authors: Tsuyoshi Hirashima, Ph.D. and Michiyuki Matsuda, M.D., Ph.D., Department of Pathology and Biology of Diseases, Graduate School of Medicine & Laboratory of Bioimaging and Cell Signaling, Graduate School of Biostudies, Kyoto University, Yoshida-Konoe-cho, Sakyo-ku, Kyoto 606-8501, Japan.

## Abstract

During collective migration of epithelial cells, the migration direction is aligned over a large, tissue-scale expanse. Although the collective cell migration is known to be directed by mechanical forces transmitted via cell-cell junctions, it remains elusive how the intercellular force transmission is coordinated with intracellular biochemical signaling to achieve collective movements. Here we show that intercellular coupling of extracellular signal-regulated kinase (ERK)-mediated mechanochemical feedback in individual cells yields long-distance transmission of guidance cues. Mechanical stretch activates ERK through epidermal growth factor receptor (EGFR) activation, and the ERK activation triggers cell contraction. In addition, the contraction of the activated cell pulls neighboring cells via cell-cell junctions, evoking another round of ERK activation and contraction in the neighbors. Furthermore, anisotropic contraction based on front-rear cell polarization guarantees unidirectional propagation of ERK activation waves, and in turn, the ERK activation waves direct multicellular alignment of the polarity, leading to long-range ordered migration. Our findings reveal that mechanical forces mediate intercellular signaling underlying sustained transmission of guidance cues for collective cell migration.

## Introduction

Collective cell migration underpins various fundamental biological processes, including embryonic development and tissue repair^1,2^. A migrating cell cluster exhibits multicellular coordination of cellular parameters, such as cytoskeleton organization^3^, organelle positioning^3-5^, and cell velocity^6,7^. Directionality of these parameters in each cell can be provided by two mechanisms. First, cells sense a direction from global external cues, such as a gradient of chemoattractant or substrate stiffness^8,9^. Second, directional cues are transmitted from a leading edge of a cell cluster to the bulk; cells located at the edge, referred to as leader cells, sense the microenvironment to spread out and dictate the direction of follower cells located behind the leader cells^10-12^. In the latter case, all follower cells as well as the leader cells generate mechanical forces to actively migrate, and those forces are orchestrated as cooperative intercellular forces over the cell cluster^13-15^. It has also been shown that the mechanical forces transmitted via cell-cell junctions provide local cues to direct an ordered cell migration in a cluster^14,16^; however, our understanding of the signaling molecules responsible for the long-range transmission of the mechanical forces is far from complete.

Extracellular signal-regulated kinase (ERK), a serine/threonine kinase, plays critical roles in mechanotransduction that regulates differentiation^17^, epithelial cell division^18^, and tissue homeostasis^19^. Earlier studies have shown that ERK activation propagates as multiple waves from leader cells to the follower cells during collective cell migration using *in vitro* cultured cells^20-22^ and *in vivo* mouse ear skin^23^. Recently, we have demonstrated that the ERK activation waves orient the directed migration of a cell cluster against the wave direction, indicating a critical role of the ERK activation waves in coordinated cell migration^24^. However, it remains largely unknown how cells harness the mechanotransduction and the ERK activation waves to coordinate collective cell migration.

In this study, by combining Förster resonance energy transfer (FRET)-based biosensors^25,26^, an optogenetic tool^27^, traction force microscopy^13^, and mathematical modeling, we show that each follower cell possesses a mechanochemical feedback system, in which stretch-induced ERK activation triggers cell contraction. Intercellular coupling of the ERK-mediated mechanochemical feedback enables sustained propagation of ERK activation and contractile force generation, leading to multicellular alignment of front-rear cell polarity over long distances. Thus, our study clarifies a mechanism of intercellular communication underlying long-range sustained transmission of directional cues for collective cell migration.

## Results

### Cell deformation waves precede ERK activation waves

To investigate the relationship between ERK activation and cell deformation during collective cell migration, Madin-Darby canine kidney (MDCK) cells confluently seeded within compartments of a silicone confinement were released for collective cell migration. ERK activity and cell deformation were evaluated by FRET imaging with EKAREV-NLS, by which ERK activation is quantified as an increase in the FRET/CFP ratio^26^, and particle image velocimetry (PIV)-based image processing, respectively. As the cells migrated toward free spaces, sustained ERK activation waves were propagated from the leader cells to the follower cells (Fig. 1a, b, Supplementary Video 1), in agreement with previous reports^20,24^. The cell deformation, i.e., extension and shrinkage, was quantified by the cell strain rate in the direction of collective cell migration (x-strain rate). Similar to the ERK activity, the x-strain rate exhibited repeated positive and negative values, namely extension and shrinkage, which propagated from the leader cells to the follower cells (Fig. 1a, c, Supplementary Video 1), as described previously^6,15^. Hereinafter, we refer to the propagation of the x-strain rate as cell deformation waves^28^.

**Fig. 1:**
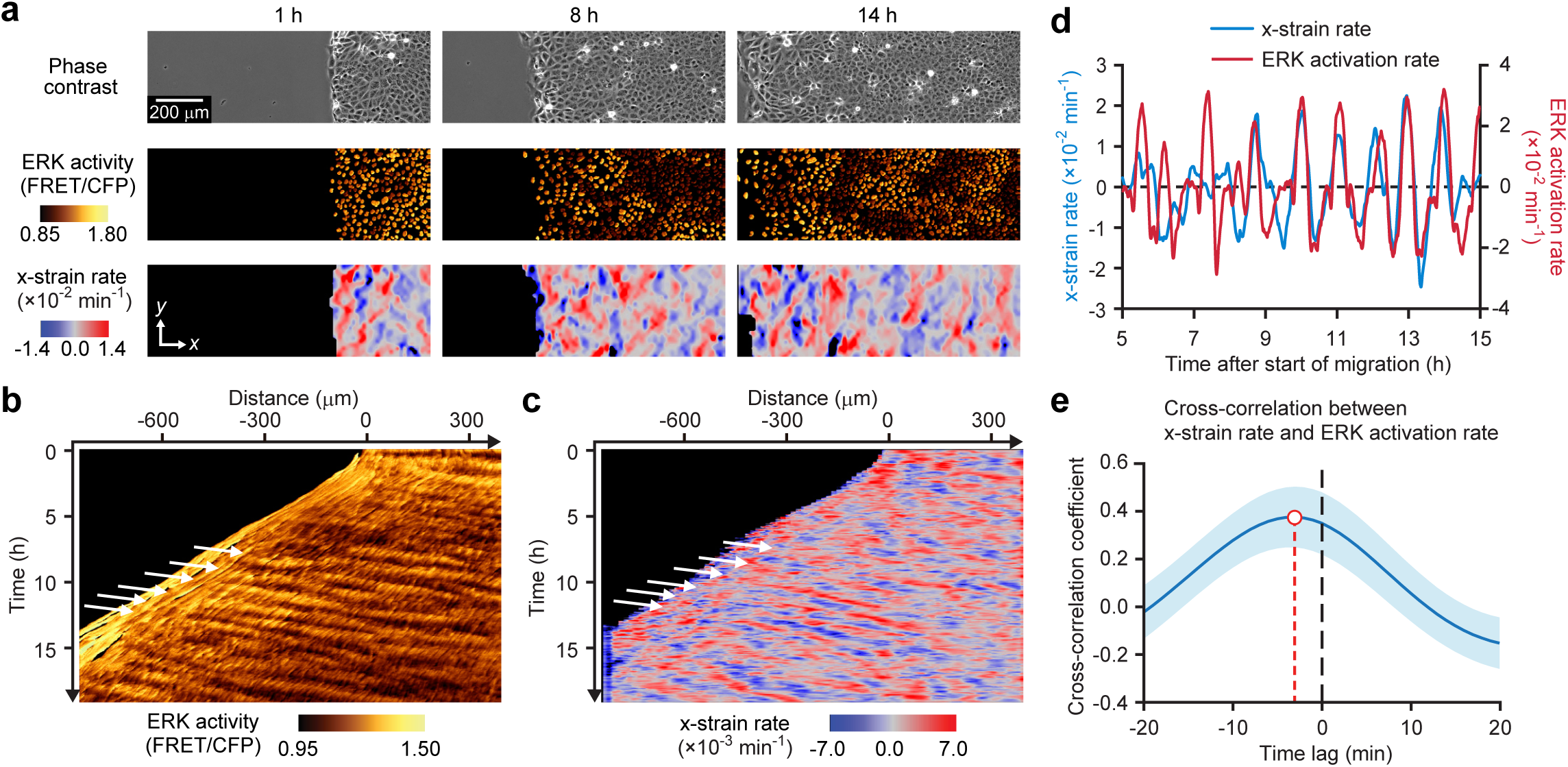
Quantitative characterization of ERK activation waves and cell deformation. **a**, Phase contrast images (upper), FRET/CFP ratio indicating ERK activity (middle), and strain rate in the direction of collective cell migration (x-strain rate; lower) are represented at 1h (left), 8 h (center), and 14 h (right) after start of time-lapse imaging. In the x-strain rate images, extending and shrinking regions are shown in red and blue, respectively. Scale bar, 200 µm. **b, c**, x-t kymographs of ERK activity (**b**) and x-strain rate (**c**), from (**a**). White arrows indicate the rightward propagation of ERK activation waves (**b**) and cell deformation waves (**c**). **d**, Temporal change of ERK activation rate and x-strain rate in a representative cell. **e**, Temporal cross-correlations between x-strain rate and ERK activation rate. The blue line indicates the average temporal cross-correlation coefficients with standard deviations (SDs). *n* = 212 cells from three independent experiments.

To analyze the correlation between the ERK activation and the cell deformation, time-series data of the ERK activity and those of the x-strain rate were collected at the single cell level. We noticed that the ERK activation rate, i.e., the ERK activity change per minute, and the x-strain rate oscillated almost synchronously with an approximately 90 min period (Fig. 1d). Furthermore, temporal cross-correlation analysis revealed that the ERK activation rate lagged 3 min behind the x-strain rate (Fig. 1e), indicating that the cell deformation waves precede the ERK activation waves. In other words, cells are first extended, followed by ERK activation, and then the cells start contracting.

### Cell extension triggers ERK activation via EGFR signaling

Because the cell extension precedes ERK activation, we reasoned that mechanical stretch activates ERK during collective cell migration. To demonstrate this, we stretched MDCK cells plated on an elastic silicone plate and compared the ERK activity before and after the mechanical stretch. As anticipated, uniaxial stretch of the MDCK cells resulted in ERK activation (Fig. 2a, b), indicating that passive extension of cells activates ERK.

**Fig. 2:**
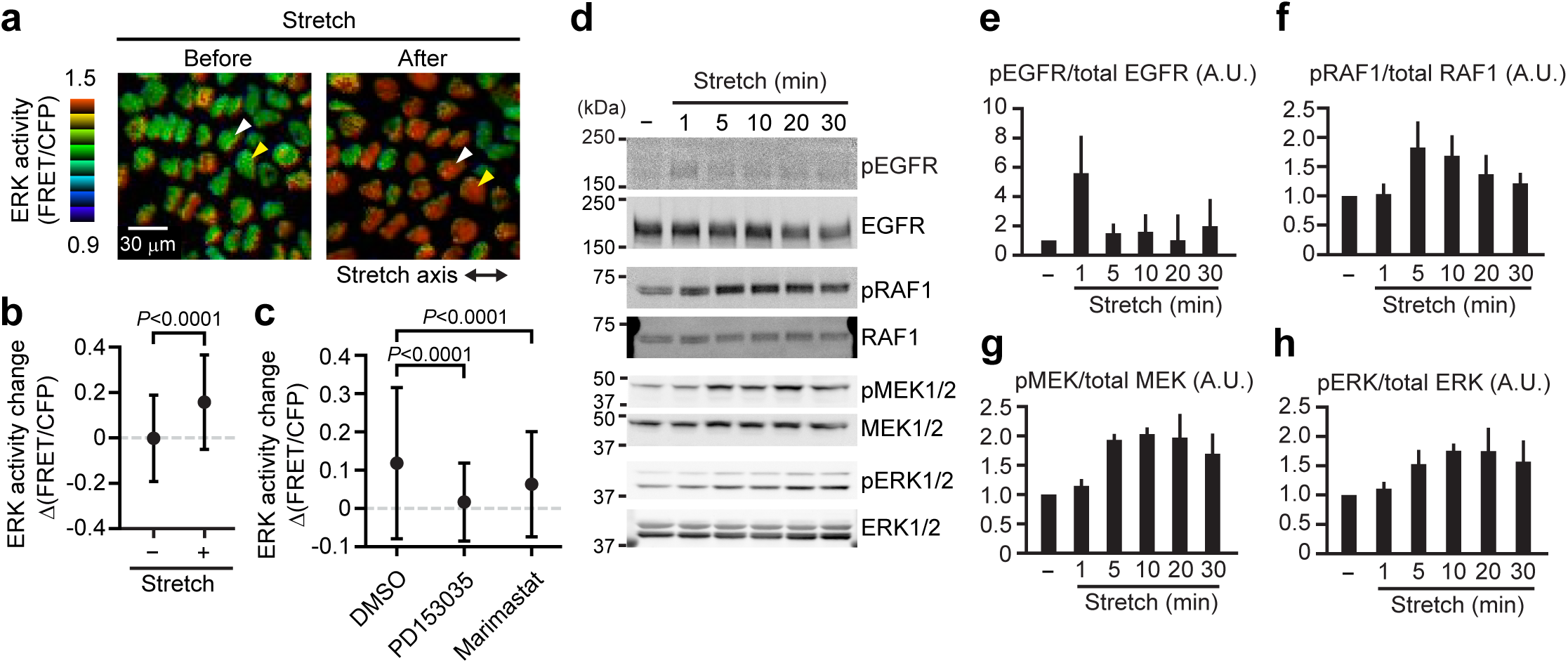
ERK activity response to mechanical perturbation. **a, b**, ERK activity change in MDCK cells on an elastic chamber. **a**, A representative image before (left) and 20 min after uniaxial stretch (right). The black double-headed arrow indicates the axis of the uniaxial stretch. White and yellow arrowheads each correspond to the same cells in the left and right images. Scale bar, 30 µm. **b**, Mean ERK activity change during 20 min without stretch and the change in ERK activity from immediately after to 20 min after the stretch, with SDs. *n* = 3612 cells (without stretch) and 3178 cells (with stretch) from three independent experiments. Unpaired *t*-test, *P* < 0.0001. **c**, ERK activity changes in cells treated with DMSO, 1 µM PD153035, and 10 µM marimastat, from immediately after to 20 min after the stretch are shown as means with SDs. *n* = 3634 cells (DMSO), 3237 cells (PD153035), and 4000 cells (marimastat) from three independent experiments. Unpaired two-tailed *t*-test, *P* < 0.0001. **d**, MDCK cells were stretched and then lysed at the indicated time points. The cell lysates were analyzed by immunoblotting with the indicated antibodies. **e**-**h**, Normalized phosphorylation levels of EGFR (**e**), RAF1 (**f**), MEK1/2 (**g**), and ERK1/2 (**h**) are represented as means with SDs (*n* = 3).

Previously, we reported that EGFR and a disintegrin and metalloprotease 17 (ADAM17), which catalyzes the shedding of membrane-tethered EGFR ligands^29^, are necessary for ERK activation waves^24,27^. Consistent with those reports, either treatment with an inhibitor of EGFR, PD153035, or that of matrix metalloproteinases (MMPs) and ADAMs, marimastat, suppressed the ERK activation waves during collective cell migration (Extended Data Fig. 1). These observations strongly suggest that the stretch-induced ERK activation requires EGFR and ADAM17 activity. In fact, both PD153035 and marimastat suppressed the stretch-induced ERK activation (Fig. 2c). Remarkably, immunoblotting showed that cell stretch transiently increased auto-phosphorylated EGFR, followed by phosphorylation of the downstream kinases including RAF1, MEK1/2, and ERK1/2 (Fig. 2d-h). Together, these results indicate that cell stretch activates EGFR and its downstream signaling molecules, including ERK.

### ERK activation induces cell contraction via Rho-associated kinase activation

Because the ERK activation precedes cell shrinkage (Fig. 1e), we speculated that ERK activation would induce cell contraction. To prove this hypothesis, we used 2paRAF, a light-inducible ERK activation system based on cryptochrome 2 (CRY2)–CIBN dimerization^27,30,31^. In this system, RAF1 fused with CRY2 is recruited to the plasma membrane by heterodimerization of CRY2 and membrane-anchored CIBN upon blue light exposure, culminating in ERK activation. We seeded cells with and without 2paRAF (2paCRY2-RAF1/CIBN-mScarlet-I-CAAX) expression into two separated compartments of a silicone confinement, respectively, and formed an interface of the two cell populations by the removal of the confinement (Fig. 3a). Upon ERK activation by blue light exposure, the interface shifted toward the side of the cells expressing 2paRAF (Fig. 3b, c, Supplementary Video 2). Moreover, inhibiting the ERK activation with the trametinib treatment suppressed the interface shift (Fig. 3b, c, Supplementary Video 2). Thus, these results clearly indicate that the ERK activation induces cell contraction of confluent MDCK cells. To confirm ERK activation also triggers cell contraction during collective cell migration, we tested the effect of ERK inhibition either with an inhibitor of EGFR, PD153035, or that of MAPK/ERK kinase (MEK), trametinib, on deformation of migrating cells. Both PD153035 and trametinib treatment damped the oscillation of the x-strain rate (Extended Data Fig. 2). Thus, ERK activation is required for cell deformation during collective cell migration.

**Fig. 3:**
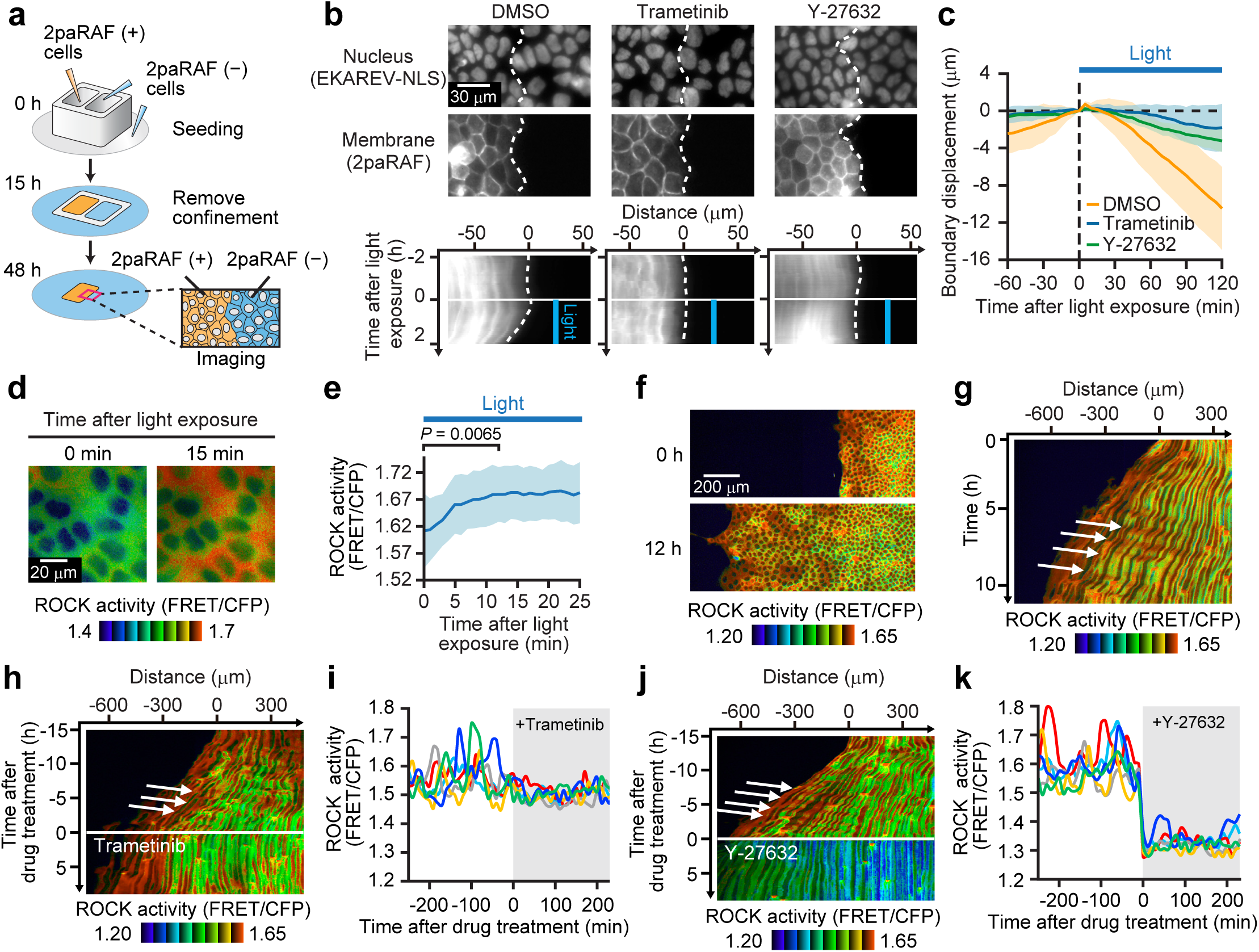
Optogenetics reveal the ERK–ROCK pathway induces cell contraction. **a**, Schematics of an experiment with the light-inducible system. The boundary between the cell populations with and without 2paRAF expression was imaged. **b**, Representative images of CFP (EKAREV-NLS; upper) and CIBN-mScarlet-I-CAAX (2paRAF; middle) treated with DMSO (left), 200 nM trametinib (center), and 10 µM Y-27632 (right) were obtained at the boundary between the cells with and without 2paRAF. Lower panels indicate kymographs of the CIBN-mScarlet-I-CAAX (2paRAF) images. Blue lines indicate the blue light illumination. Scale bar, 30 µm. **c**, Displacement of the boundary is plotted over time after blue light exposure. The lines represent the average with SDs. *n* = 9 from three independent experiments. **d**, ROCK activity images of MDCK cells expressing 2paRAF and a ROCK biosensor are represented at 0 min (left) and 15 min (right) after the start of blue light exposure. Scale bar, 20 µm. **e**, Quantification of ROCK activity in each cell in (**d**) after the start of blue light exposure. The line represents the average with SDs. *n* = 15 from three independent experiments. Unpaired *t*-test, *P* = 0.0065. **f**, ROCK activity images are represented at 0 h (upper) and 12 h (lower) after removal of confinement. Scale bar, 200 µm. **g**, A kymograph of the ROCK activity in (**f**). White arrows indicate the rightward propagation of ROCK activity. **h, j**, A kymograph of ROCK activity is shown before and after 200 nM trametinib (**h**) and 100 µM Y-27632 (**j**) treatment. **i, k**, ROCK activity in 5 representative cells was plotted over time after trametinib (**i**) and Y-27632 (**k**) treatment.

Cell contraction is driven by actomyosin, which involves Rho-associated kinase (ROCK)-mediated regulation^32-34^. Thus, we examined the effects of a ROCK inhibitor, Y-27632, on the ERK-induced cell contraction. We found that Y-27632 suppressed the ERK-induced cell contraction (Fig. 3b, c, Supplementary Video 2). This observation prompted us to examine whether ERK induces ROCK activation. To this end, we used a cytosolic FRET biosensor for ROCK activity^35^. After optogenetic ERK activation, ROCK activity increased, then plateaued within 12 min after the blue light exposure (Fig. 3d, e). We next examined the ROCK activity dynamics during collective cell migration. Interestingly, ROCK activity exhibited repeated unidirectional wave propagation from leader cells to the follower cells, as did the ERK activity (Fig. 3f, g, Supplementary Video 3). Inhibition of ERK with trametinib abolished the propagation of ROCK activation waves (Fig. 3h, Supplementary Video 3) as well as temporal oscillatory activation in each cell (Fig. 3i). In stark contrast, inhibition of ROCK with Y-27632 abolished the oscillatory activity in each cell and also decreased the basal activity of ROCK drastically (Fig. 3j, k, Supplementary Video 3), indicating that ERK activity is responsible only for the oscillatory component of ROCK activity. Thus, we conclude that ROCK is activated downstream of ERK and is integral to ERK-induced cell contraction.

### ERK activation decreases traction forces and accumulates F-actin at the cell-cell interface

How does ERK activation alter mechanical force generation to induce contraction? To answer this question, we combined light-inducible ERK activation with traction force microscopy^13^, which allows the measurement of traction forces loaded by cells on the substrate. On a substrate of polyacrylamide gel embedded with fluorescent beads, the interface between the cells with and without 2paRAF expression was formed, as already described (Fig. 3a). The traction force exerted by the 2paRAF-expressing cells markedly decreased within 40 min after the start of blue light exposure (Fig. 4a, b, Supplementary Video 4). By contrast, inhibiting ERK with trametinib restored the traction force generation (Fig. 4c), confirming that ERK activation suppresses force loading on the substrate. The gradual increase in traction force by the cells without 2paRAF expression in Fig. 4A, B is independent of the optogenetic ERK activation in the neighboring 2paRAF-expressing cell cluster because the traction force increased over time even without blue light exposure (Extended Data Fig. 3). This is likely because increasing cell density by proliferation downregulates ERK activity^27^. Moreover, we found that subcellular localization of F-actin in migrating cells is altered depending on the ERK activity level (Fig. 4d, e). ERK activation by epidermal growth factor (EGF) promoted F-actin localization at the lateral side of the cells. By contrast, ERK inhibition by trametinib treatment produced stress fibers at the basal side of the cells. Therefore, these results suggest that ERK activation triggers cell contraction by accumulating F-actin at the cell-cell interface and triggering predominant force loading on the interface while suppressing traction force generation.

**Fig. 4:**
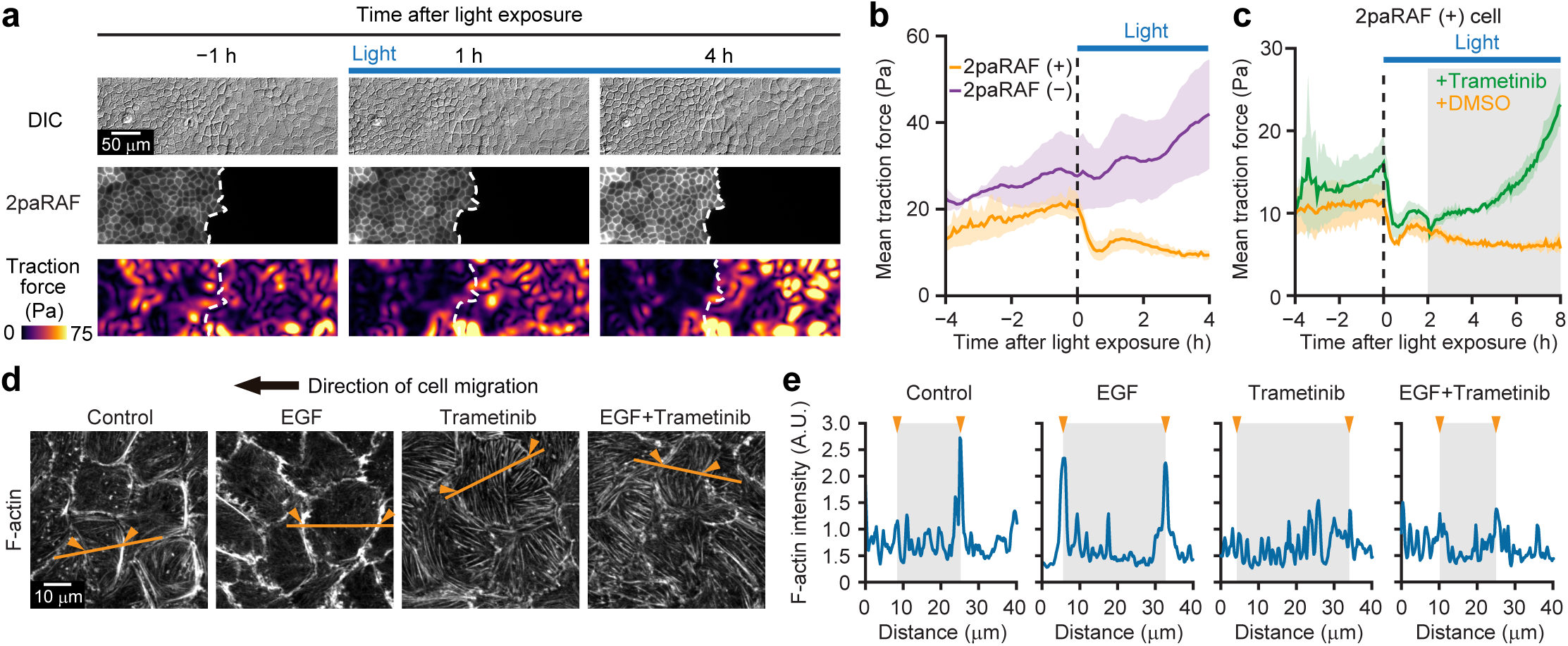
ERK activation decreases the traction force and accumulates F-actin at the cell-cell interface. **a**, Traction force microscopy with optogenetic ERK activation. Differential interference contrast (DIC) images (upper), CIBN-mScarlet-I-CAAX fluorescence (2paRAF; middle), and traction force (lower) are represented at -1 h (left), 1 h (center), and 4 h after the start of blue light exposure. Scale bar, 50 µm. **b**, Mean traction force in cells with and without 2paRAF, as shown in (**a**), with SDs (n = 3). **c**, Mean traction force in cells with 2paRAF with DMSO or 200 nM trametinib treatment at 2 h after the start of blue light exposure (n = 3). **d**, Fluorescence images of F-actin (phalloidin) in migrating MDCK cells 45 min after treatment with DMSO, 100 ng mL^−1^ EGF, 200 nM trametinib, or both EGF and trametinib. Scale bar, 10 µm. **e**, Intensity profile of F-actin along the orange lines in (**d**). Orange arrowheads indicate edges of cells.

### Zonal cell contraction initiates sustained unidirectional ERK activation waves at the interface

We have shown that each cell possesses an ERK-mediated mechanochemical feedback system coupling cell deformation and ERK activation: ERK is activated by cell extension, and the activated ERK induces cell contraction. Considering the tight physical connection between epithelial cells, contraction of a cell cluster should stretch cells in the adjacent cluster, mainly at its border, thereby generating the ERK activity propagation. To test this hypothesis, we used a rapamycin-activatable (RA) Rho guanine nucleotide exchange factor (GEF) system to induce contraction of a cell cluster^36,37^. We first seeded cells carrying the RA Rho GEF expression system into a confinement (Fig. 5a), and then seeded the cells with the EKAREV-NLS after the removal of the confinement, resulting in the formation of an interface between cells expressing RA Rho GEF and cells expressing EKAREV-NLS. Before rapamycin treatment, ERK activity was randomly propagated without any preferential direction (Fig. 5b, Supplementary Video 5). Upon rapamycin treatment, the cells with RA Rho GEF began to contract, resulting in the extension of the adjacent EKAREV-NLS–expressing cell cluster toward the RA Rho GEF-expressing cell clusters (Fig. 5c, d, Supplementary Video 5). With this trigger, ERK activation waves emerged at the interface and were unidirectionally propagated toward the extended EKAREV-NLS–expressing cell cluster (Fig. 5c-e). In addition, the displacement of the EKAREV-NLS–expressing cells was oriented toward the RA Rho GEF-expressing cell clusters and was directed opposite the ERK activation waves (Fig. 5f). Thus, cell contraction generates ERK activation waves in the adjacent cell cluster. We then asked whether a cell-cell tight connection is required for intercellular propagation of ERK activation waves. To test this, we knocked-out α-catenin, a major cell-cell junction component^38^, in MDCK cells (Fig. 5g). As expected, E-cadherin at the cell-cell junction was reduced in the α-catenin KO cells in comparison with wild-type (WT) cells (Fig. 5h). Coordinated ERK activity propagation was severely disrupted in the α-catenin KO cells (Fig. 5i, j, Supplementary Video 6), indicating that intercellular mechanical linkage is required for ERK activity propagation. Thus, we conclude that intercellular force transmission mediates ERK activation waves.

**Fig. 5:**
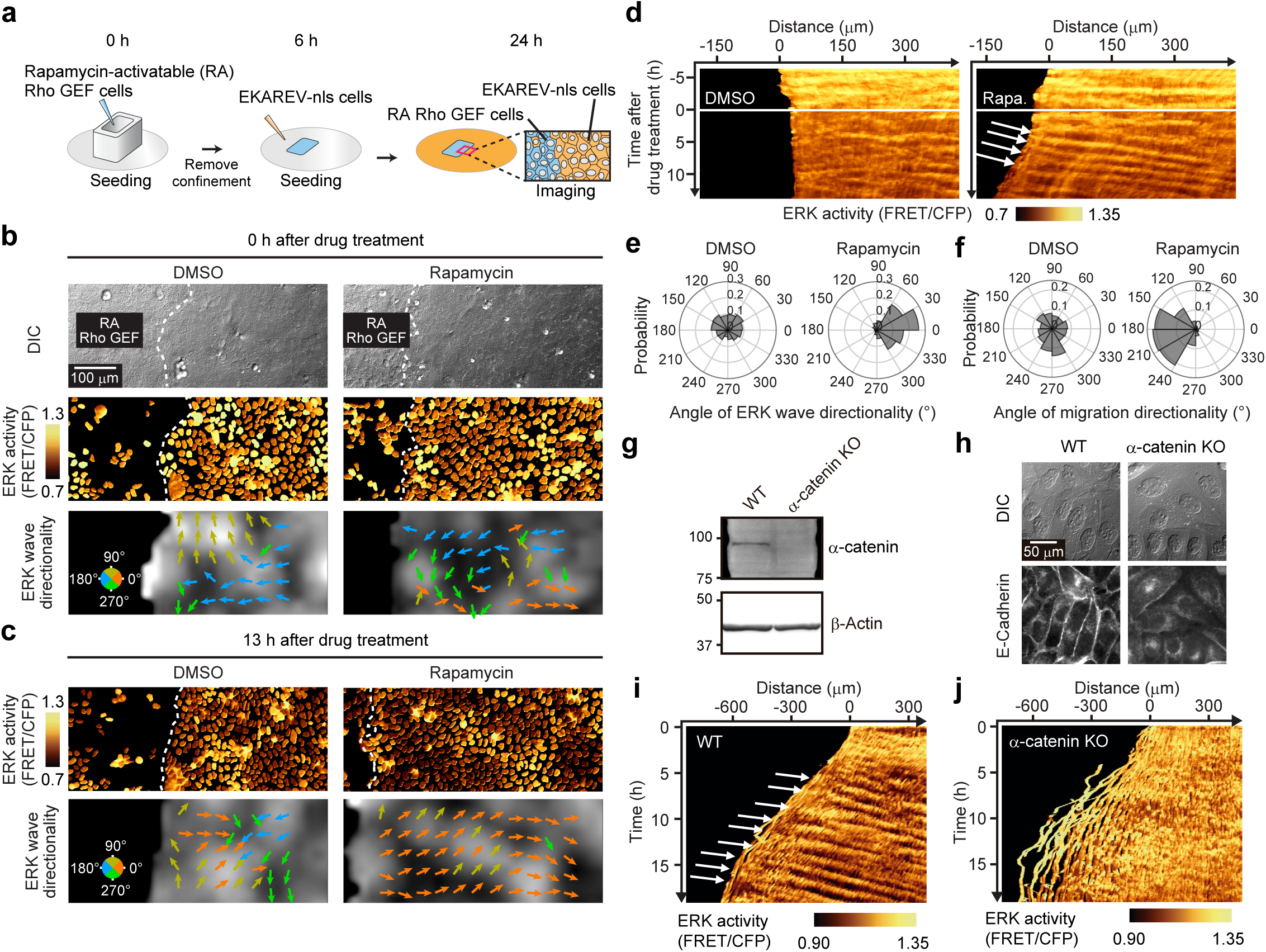
ERK activation waves are mediated by intercellular mechanical force. **a**, Schematics of an experiment using the rapamycin-inducible system. The boundary between the cell population with RA Rho GEF and that with EKAREV-NLS expression was imaged. **b**, DIC images (upper), ERK activity (middle), and ERK wave directionality (lower) are represented at 0 h after DMSO (left) and 50 nM rapamycin (right) treatment. Broken lines indicate the cell population boundary. Arrows represent the direction of ERK activity propagation. For the analysis of ERK wave directionality, the EKAREV-NLS–expressing cells located to the left of the cell population boundary were omitted from the analysis. Scale bar, 100 µm. **c**, ERK activity (upper) and ERK wave directionality (lower) are represented at 13 h after the drug treatment. **d**, Kymographs of ERK activity with DMSO (left) or rapamycin (right) treatment. The EKAREV-NLS–expressing cells located on the left side of the cell population boundary were excluded from the kymographs for visibility. **e, f**, Polar histograms showing the distribution of ERK wave (**e**) and cell displacement (**f**) direction over 13-13.5 h after treatment with DMSO (left) and rapamycin (right). For ERK wave direction, *n* = 1078 (DMSO) and 1127 (rapamycin) from three independent experiments. For cell displacement direction, *n* = 475 (DMSO) and 1167 (rapamycin) from three independent experiments. **g**, Analysis of the expression level of α-catenin and β-actin in WT and α-catenin KO MDCK cells by immunoblotting. **h**, DIC images (upper) and immunofluorescence images of E-cadherin (lower) in the confluent WT and α-catenin KO MDCK cells. **i, j**, Kymographs of ERK activity during cell migration in WT (**i**) and α-catenin KO (**j**) MDCK cells.

### Front-rear cell polarization is required for the unidirectional propagation of ERK activation waves

We next investigated how unidirectional propagation of ERK activation waves is achieved during collective cell migration. If cells contract isotropically, the loss of directional information would impede the unidirectional propagation of ERK activation waves. Therefore, there must be a mechanism by which the directional information is retained during the wave propagation. We found that the cell extends preferentially along the direction of migration (Fig. 6a, b), indicating that cells deform anisotropically during collective cell migration. Furthermore, phosphorylated myosin light chain, a marker of contractile myosin, and F-actin accumulated at the basal rear side of migrating cells (Fig. 6c, Extended Data Fig. 4a, b), which are indicative of polarized rear contraction. The presence of a front-rear cell polarity was also confirmed by the localization of GM130, a Golgi apparatus marker, known to be located in front of the nucleus in some types of migrating cells^39,40^. When cells were examined immediately after release from confinement (0 h migration), the Golgi orientation relative to the nucleus was not biased toward the leading edge (Extended Data Fig. 4c, d). By contrast, 21 h after the initiation of collective cell migration, the Golgi orientation was biased toward the direction of the migration over more than a 1 mm range from the leader cells (Fig. 6d, e). To quantify the degree of alignment, we defined the directedness of the Golgi orientation (Fig. 6f), and found that the Golgi directedness first increased around the leader cells, and then the increase spread through the follower cells over time (Fig. 6g). In addition, the directedness of ERK activation waves and that of migration showed similar dynamics to the Golgi directedness (Fig. 6h, i). Thus, these results clarify that alignment of front-rear cell polarity in follower cells positively correlates with unidirectionality of ERK activation waves.

**Fig. 6:**
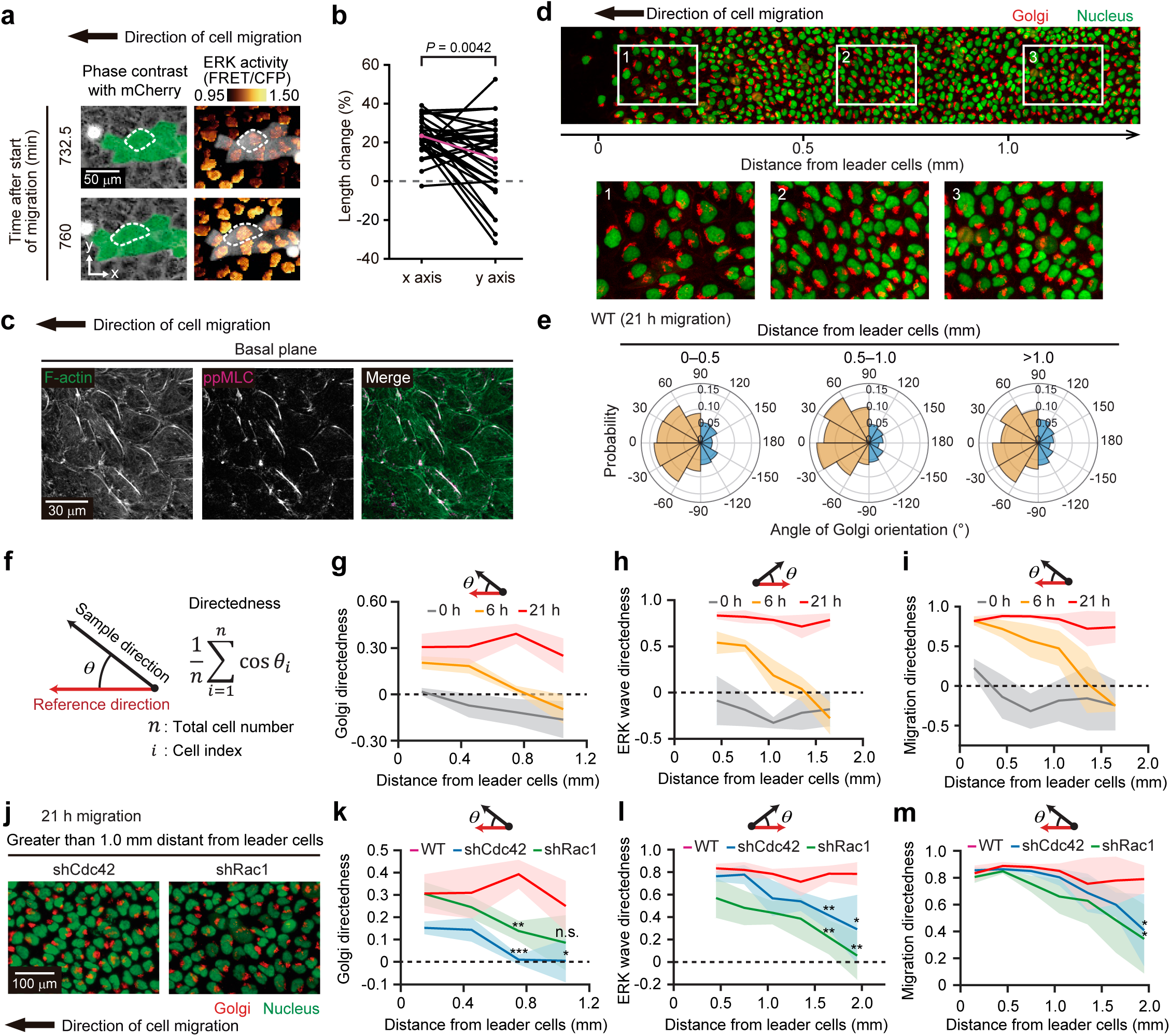
Multicellular alignment of front-rear polarization underpins unidirectional ERK activation waves. **a**, A representative cell expressing mCherry in the cytosol is marked by a broken line in contracted (upper) and extended (lower) states during collective cell migration. The left column shows composite images of phase contrast and mCherry fluorescence (green), and the right column shows ERK activity. Cells showed low ERK activity in the contracted state and high ERK activity in the extended state. **b**, Changes in lengths of cells along the x axis and y axis between the contracted and extended states are plotted. Each line indicates an individual cell. *N* = 30 cells from three independent experiments. Paired *t*-test, *P* = 0.0042. **c**, Immunofluorescence images of F-actin (left) and di-phosphorylated MLC (ppMLC; center) at the basal plane in MDCK cells at 17 h after migration. The right column indicates the composite images of F-actin and ppMLC. **d**, Immunofluorescence images of the Golgi apparatus (GM130) and the nucleus (EKAREV-NLS) in MDCK cells at 21 h after migration. The upper image shows a wide-view field, and the lower images are magnified images of the regions corresponding to the numbered windows in the upper image. **e**, Polar histograms showing the distribution of Golgi orientation relative to the nucleus in the cells at 0-0.5 mm (left), 0.5-1.0 (center), and greater than 1.0 mm (right) distant from the leader cells. *n* = 1297 cells (0-0.5 mm), 2096 cells (0.5-1.0 mm), and 1452 cells (>1.0 mm) from three independent experiments. **f**, θ is the angle between the reference direction and sample direction. Directedness was calculated by the indicated equation. The reference direction is set to the left for analysis of Golgi orientation and cell migration direction, and to the right for ERK wave direction, in order to match the signs of their directedness. **g**-**i**, Mean directedness of Golgi orientation (**g**), ERK wave direction (**h**), and cell migration (**i**) binned every 300 µm from the leader cells after 0 h, 6 h, and 21 h migration is plotted over the distance from the leader cells, with SDs. The data were obtained from three independent experiments. The first bin of the ERK wave directedness was excluded from the result because of outliers due to boundary effects. **j**, Immunofluorescence images of Golgi (GM130) and nuclei (EKAREV-NLS) in shRac1- and shCdc42-expressing MDCK cells at regions greater than 1 mm from the leader cells after 21 h migration. **k**-**m**, Directedness of Golgi orientation (unpaired *t*-test; \*\*\**P* = 0.0006 (0.6-0.9 µm, WT versus shCdc42), \**P* = 0.0456 (0.9-1.2 mm, WT versus shCdc42), \*\**P* = 0.0051 (0.6-0.9 µm, WT versus shRac1)) (**k**), ERK wave direction (unpaired *t*-test, \*\**P* = 0.0053 (1.5-1.8 mm, WT versus shCdc42), \**P* = 0.0378 (1.8-2.1 mm, WT versus shCdc42), \*\**P* = 0.0011 (1.5-1.8 mm, WT versus shRac1), \*\**P* = 0.0018 (1.8-2.1 mm, WT versus shRac1)) (**l**), and cell migration direction (unpaired *t*-test, \**P* = 0.0183 (1.8-2.1 mm, WT versus shRac1), \**P* = 0.0374 (1.8-2.1 mm, WT versus shCdc42)) (**m**) binned every 300 µm from the leader cells after 21 h migration were plotted over the distance from the leader cells with SDs. The data of WT cells is the same as that of 21 h migration in (**g**-**i**). The data were obtained from three independent experiments.

To investigate the requirement of front-rear cell polarization for the unidirectional ERK activation waves, we knocked down Rac1 or Cdc42, which are required for the establishment of front-rear polarization^39^. Expression of short hairpin RNAs reduced Rac1 and Cdc42 expression to 50% and 30%, respectively (Extended Data Fig. 4e, f). In the knocked-down cells, the front-rear polarization of Golgi in the leader cells was still directed in the direction of the migration; however, that of the follower cells was impaired after 21 h migration (Fig. 6j, k). Moreover, the unidirectionality of the ERK activation waves and the alignment of migration direction were also impaired in the follower cells of the knocked-down cells (Fig. 6l, m, Supplementary Video 7). Taken together, these results demonstrate that the alignment of front-rear polarization toward the direction of the migration is required for the unidirectional ERK activation waves and the ordered cell migration over long distances from the leading edge.

### ERK activation waves orient front-rear polarity in follower cells

We further addressed the relationship between the ERK activation waves and the front-rear polarization. Previously, it has been proposed that force loading on cell-cell junctions directs front-rear polarization^16^. Consistent with this, disruption of cell-cell junctions by α-catenin KO abolished the alignment of the front-rear polarization toward the direction of migration (Extended Data Fig. 5a-c). From this observation, we speculated that ERK-induced contractile force generation would direct the front-rear polarization in neighboring follower cells. As expected, inhibition of ERK activation with trametinib suppressed the alignment of the front-rear polarization and migration directionality in the follower cells at 21 h after migration (Fig. 7a-c, Extended Data Fig. 5d, Supplementary Video 8). Interestingly, constitutive ERK activation by 12-*O*-tetradecanoylphorbol 13-acetate (TPA) also inhibited the alignment of the front-rear polarization and migration directionality in the follower cells (Fig. 7a-c, Extended Data Fig. 5e, Supplementary Video 8), suggesting that periodic ERK activation in the form of waves is important for polarizing follower cells. We then tested whether synthetic ERK activation waves can enhance the directedness of the Golgi polarization in the follower cells. To this end, we used MDCK cells expressing 2paRAF (2paCRY2-RAF1/CIBN-CAAX) and Golgi-7-mCherry as a Golgi apparatus marker. Those cells were confluently seeded in a confinement, and then released for migration with an EGFR inhibitor PD153035 to suppress autonomous ERK activation waves (Fig. 7d). Of note, inhibition of EGFR signaling suppresses ERK activation waves in follower cells, but ERK activation in the cells near the leading edge remains (Extended Data Fig. 1b, c). Thus, the leader cells still migrate, even with EGFR signaling inhibition, while follower cell migration is abbrogated^24^. Under this condition, we created optogenetic ERK activation waves by shifting patterned blue light-illumination to mimic spontaneous ERK activation waves; i.e., ∼35 µm width and 400 µm intervals at 3 µm min^-1^ velocity (Fig. 7d). With the synthetic ERK activation waves, the front-rear cell polarizations were significantly aligned even in follower cells more than 600 µm distant from the leader cells, compared with the samples in the absence of ERK activation waves (Fig. 7e, f, Supplementary Video 9). Thus, these results clearly demonstrate that ERK activation waves orient the front-rear polarity in cells against the direction of the waves. Collectively, we conclude that unidirectional ERK activation waves and multicellular alignment of the front-rear polarization propagate cooperatively, enabling long-distance transmission of directional cues for collective cell migration.

**Fig. 7:**
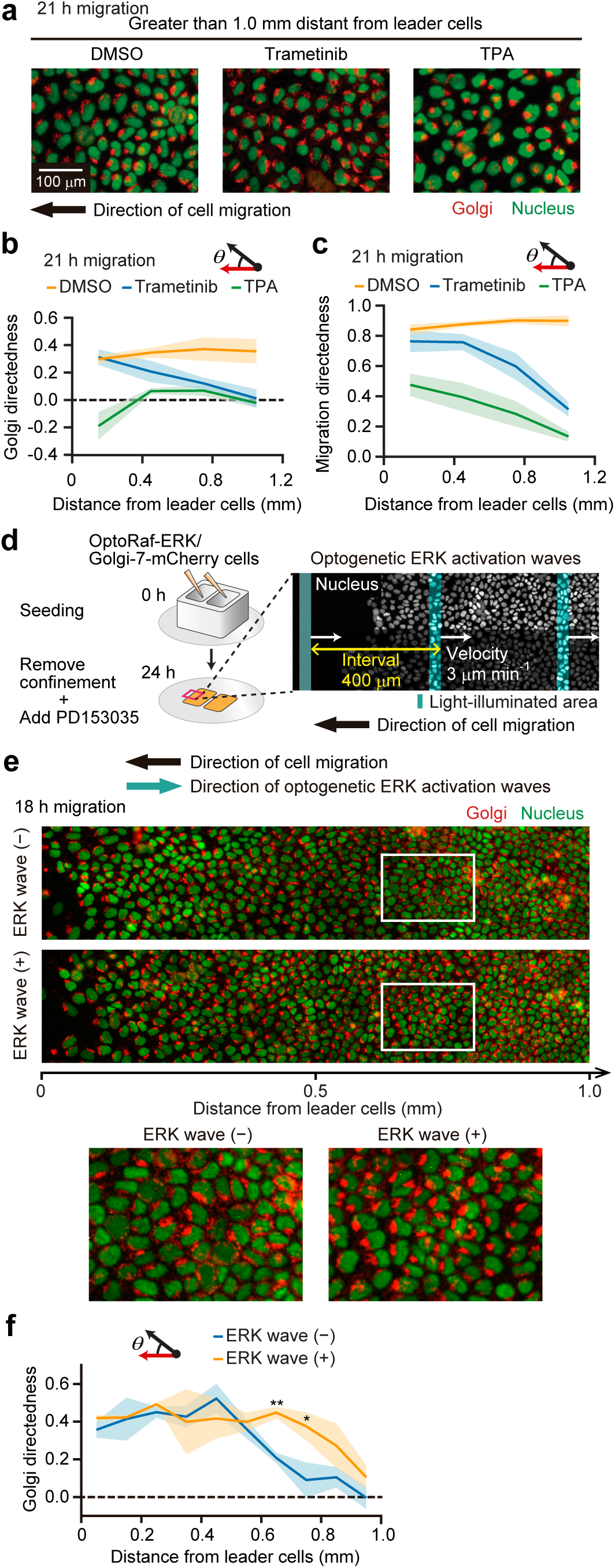
Synthetic ERK activation waves induce polarity ordering. **a**, Immunofluorescence images of Golgi (GM130) and nuclei (EKAREV-NLS) in MDCK cells treated with DMSO (left), 200 nM trametinib (center), and 10 nM TPA (right) at a region more than 1.0 mm distant from the leader cells at 21 h after migration. Scale bar, 100 µm. **b, c**, Mean directedness of Golgi orientation (**b**) and cell migration direction (**c**) binned every 300 µm from the leader cells after 21 h migration were plotted over distance from the leader cells, with SDs. The data were obtained from three independent experiments. **d**, Flow diagram of an experiment on synthetic ERK activation waves. **e**, Fluorescence images of Golgi apparatus (Golgi-7-mCherry) and nuclei (EKAREV-NLS) in MDCK cells with or without optogenetic ERK activation waves for 18 h. The upper images are wide-view fields, and the lower images are magnified views of the regions outlined by white windows in the upper images. **f**, The mean directedness of Golgi orientation binned every 100 µm from the leader cells in (**e**) was plotted over distance from the leader cells, with SDs. The data were obtained from three independent experiments. Unpaired *t*-test; \*\**P* = 0.0011 (0.6-0.7 µm, plus versus minus), \**P* = 0.0267 (0.7-0.8 mm, plus versus minus).

### Modeling cellular mechanochemical feedback with polarity demonstrates long-range unidirectional ERK activation waves

Finally, we developed a mathematical model to understand the role of ERK-mediated mechanochemical waves in collective cell migration. We employed a two-dimensional cellular Potts model (CPM) to represent behavior of the epithelial monolayer at a single cell scale^41-43^. In this model, each cell morphology is represented as a cluster of regular lattices, and a state of lattices determine an energy of the system. The dynamics of a multicellular system proceeds stochastically on the basis of an energy minimization using a Monte Carlo simulation algorithm. We here regard a unit of trials in Monte Carlo simulation (Monte Carlo steps: MCS) as 1 h in experiments. The energy in the model includes essential properties in multicellular mechanics, i.e., intercellular adhesion, cell elasticity, and active cellular contraction^44,45^. In addition, the cells possess an inherent orientation regarding the polarized rear contraction, equivalent to the front-rear polarity defined as the Golgi orientation in our experiments (Fig. 8a). In our model, orientations of the cell polarity in individual cells tend to be aligned as a result of the physical interaction with neighboring cells (Supplementary notes)^46-48^. The anisotropy along the front-rear polarity and the strength of cell contraction are governed by the cell polarity parameter ω and the ERK activity, respectively. That is, the cell contraction is isotropic when ω=0 and polarized for the rear side of the cell with increasing ω (Fig. 8b); we set ω=1 as suggested in experiments, unless otherwise noted. In simulations, we assumed that the follower cells obey the stretch-induced contraction, while the leader cells keep migrating towards the free space.

**Fig. 8:**
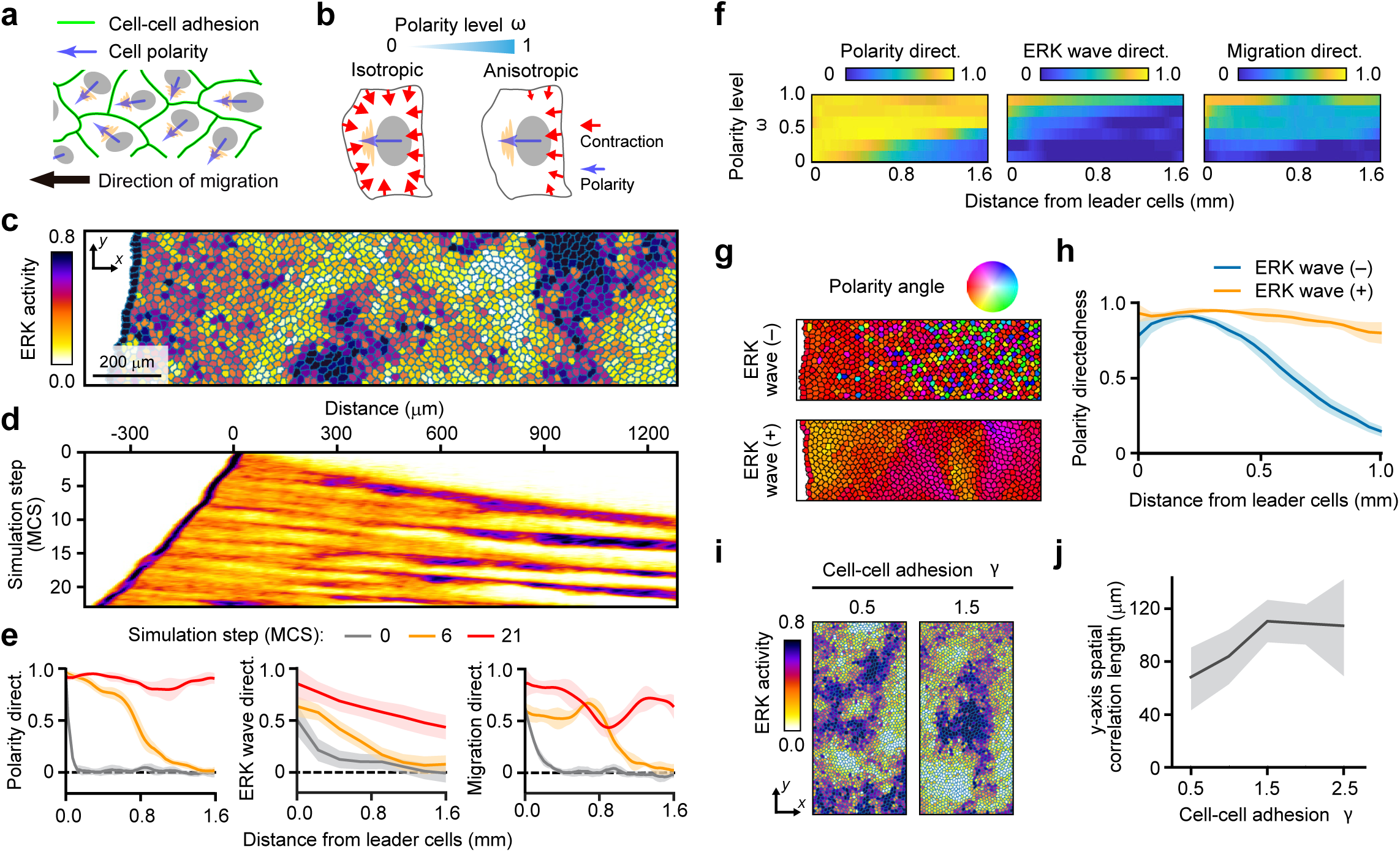
Model simulation for collective cell migration. **a**, Schematics of the mathematical model. **b**, Contraction mode depending on the parameter ω, defined as the polarity level. **c**, Snapshot of ERK activity in the CPM simulation. Scale bar, 200 µm. **d**, Kymograph of ERK activity along the x-axis and simulation step. The color corresponds to values of the color bar in (**c**). **e**, Spatial distribution of mean directedness, with SDs. The color shows different simulation steps. *n*=5. **f**, The effect of polarity level ω on the directedness. The color represents mean values of the directedness. *n*=5. **g, h**, Cell polarity directedness with (+) or without (–) ERK wave. The color in (**g**) represents the angle of cell polarity. (**h**) Mean polarity directedness from the leading edge, with SDs. *n*=5. **i**, Snapshot of ERK activity distribution in weak (*γ*=0.5) and strong (*γ*=1.5) multicellular integrity. **j**, Mean y-axis spatial correlation length of the ERK activity on the intercellular integrity *γ*, with SDs. *n*=5.

We performed simulation analysis under a similar condition to the confinement release assay, and confirmed that the unidirectional ERK activation waves from the leader cells to the follower cells were reproduced (Fig. 8c, d, Supplementary Video 10). Moreover, the simulation results well mimicked experimental measurements in terms of spatio-temporal profiles of front-rear polarity directedness, ERK wave directedness, and migration directedness (Fig. 8e). We then examined the effect of the cell polarity level ω on the directedness. For any type of directedness, the spatial range with high values becomes more limited around the leading cells, with the decrease of ω (Fig. 8f, Supplementary Video 10) as observed with disruption of polarity proteins (Fig. 6j-m). This indicates that the cell anisotropy in contraction along the front-rear polarity is essential for long-range multicellular guidance. In addition, we investigated the effect of ERK activation waves on the polarity directedness (Fig. 8g and h). Even without the ERK activation waves, the front-rear polarity in the follower cells around the leading edges is relatively directed toward the front because the migrating leader cells pull near-follower cells; however, the polarity directedness gradually decreases with distance from the leader cells. By contrast, the polarity directedness is maintained long-range from the leader cells via the ERK activation waves (Fig. 8h). Hence, ERK activation waves direct the front-rear polarity of the follower cells, consistent with the experiments with optogenetic ERK activation waves (Fig. 7f).

Further in silico analysis led to the finding that the strength of cell-cell adhesion γ controls synchronization of the ERK activity along a perpendicular axis to its propagation direction (y-axis in Fig. 8i). We introduced a spatial correlation length of the ERK activity along the y-axis as a measure of its synchronization, and examined a dependency of γ on the spatial correlation length. Our numerical investigation clearly shows that the spatial correlation length increases with increasing cell-cell adhesion strength γ (Fig. 8j), indicating that multicellular integrity via intercellular mechanical linkages plays a key role in the ERK activity synchronization. Taken together, tight connections between cells, each of which experiences mechanochemical feedback with front-rear polarity, effect well-ordered long-range ERK activation waves.

## Discussion

During collective cell migration of epithelial cells, not only leader cells but also follower cells exert traction forces on their substrates to drive cell movement, resulting in long-distance coordinated cell migration^13^. Thus, the follower cells can sense the direction of the leader cell migration. Previous studies have revealed that the directional cues are transmitted to the follower cells by intercellular mechanical forces^15^. The mechanical tension varies dynamically and spreads over long distances, leading to establishment of local anisotropic stress, along which the cells tend to migrate^14,16^. However, the molecular mechanism by which the mechanical forces are sustainably transmitted over long distances to direct collective cell migration remains to be revealed. Here we identified that ERK is a key molecule regulating cellular mechanochemical feedback, which governs long-distance sustained propagation of the directional cues. We propose a mechanism of ERK and mechanical force-mediated intercellular communication underlying the collective cell migration (Extended Data Fig. 6). In the initial process of collective cell migration, advancement of the leader cells toward the free space exerts pulling forces on the adjacent follower cells, directing front-rear cell polarization. The pulling force then stretches the follower cells and activates the EGFR-ERK signaling cascade, which in turn generates contractile forces at the rear side of the follower cells through the ROCK pathway. The contraction of the follower cells not only provides the driving force of the follower cell migration, but also exerts pulling forces on the next follower cells, evoking another round of cell stretch, front-rear polarization, and ERK activation. Thus, intercellular coupling of the ERK-mediated mechanochemical feedback via cell-cell junctions enables sustained propagation of pulling forces, front-rear cell polarity, and ERK activation over a large tissue scale, leading to collective cell migration.

The combination of optogenetic ERK activation and traction force measurement revealed that ERK plays a critical role in translocating cellular force generators. Our experiments demonstrate that the ERK activation triggers cell contraction through ROCK activation (Fig. 3b-e) while it decreases the traction forces (Fig. 4a, b). The effect of ERK activation on the force generation can be explained by the translocation of dense F-actin fibers from cell bases to cell-cell junctions (Fig. 4d, e), which should lead to an efficient tug on adjacent cells. It is plausible that the ERK-induced cell contraction through ROCK involves Rho GEF activation localized at cell-cell junctions. Along this line of reasoning, a previous study demonstrated that mechanical stretching of cells caused localization of Rho GEF at cell-cell junctions, giving rise to local activation of RhoA, and thereby the junction-associated actomyosin^49^. On the other hand, the decrease in traction forces by ERK activation could be attributed to disruption of focal adhesion-associated actin stress fibers due to promotion of focal adhesion turnover^50,51^. The focal adhesion turnover should increase deformability of cells to efficiently contract upon contractile force generation.

Remarkably, our results imply that ERK activation triggers polarized rear contraction in each cell. As the intracellular ERK activation may not be significantly polarized due to rapid ERK diffusion within the cytoplasm^52^, the effect of ERK activation on the polarized rear contraction is explained by an antagonistic activation of Rac1 and RhoA^53^. Force loading on a cell-cell junction activates Rac1 via dissociation of merlin from the junction, which directs the cell front^16^. The preoccupied Rac1 activation at the cell front should restrict the activation of RhoA-regulated contractile machineries to the rear side of the cells, leading to polarized rear contraction upon ERK activation. Therefore, ERK should convey the signal to stimulate the contractile machineries within a cell at the back in response to pulling forces at the front.

An important unsolved question is how the mechanical stimuli are converted into signal transduction to activate ERK. We have previously demonstrated that ADAM17 is indispensable for the propagation of the ERK activation in MDCK cells^24^. It has been proposed that ADAM17 catalyzes the ectodomain shedding of EGFR ligands^54^, and the released ligands can bind and activate EGFR in an autocrine or paracrine manner. In this study, we found that stretch of MDCK cells activates EGFR (Fig. 2d,e), as reported in other cell types^55,56^. Therefore, we suppose that the stretch-induced EGFR activation involves the ADAM17-mediated shedding of EGFR ligands. It has been shown that ERK activates ADAM 17 by means of phosphorylation of Thr735^57-59^. However, EGFR activation occurs as early as 1 min after stretching and precedes ERK activation (Fig. 2d-h), which is at odds with the involvement of ERK activation in the stretch-induced EGFR activation. It has also been reported that src, a protein tyrosine kinase, is involved in ADAM17 activation^60,61^ and is required for the stretch-induced EGFR activation^56^. Thus, future studies will be necessary to explore the possibility of src-mediated ADAM17 activation in the ERK activation waves.

In conclusion, we have revealed that mechanical force transmission functions as a mediator of the intercellular ERK signal transduction underlying collective cell migration. Current understanding of the intercellular signal transduction mostly emphasizes the importance of intercellular transfer of biochemical molecules including growth factors, hormones, and neurotransmitters. Thus, our present study raises another consideration, which is the critical role of cellular response to mechanical stimuli in intercellular signal transduction.

## Materials and Methods

### Plasmids

pT2A-EKAREV-NLS and plasmids for RA Rho GEF (pCX4puro-LDR, and pCX4bsr-3HA-FKBP-p63RhoGEF-DH) were described previously^26,35,36,62,63^. pT2ADWpuro_2paRAF (Addgene plasmid: no. 129653) encoding 2paCRY2-RAF1-P2A-CIBN-mScarlet-I-CAAX was described previously^30^. pT2ADWpuro_2paRAFΔmScarlet-I encoding 2paCRY2-RAF1-P2A-CIBN-CAAX was generated by PCR and subcloned into the pT2ADWpuro vector. The FRET biosensor for ROCK activity, Eevee-ROCK-NES, was described previously^35^ and subcloned into a pCSII vector. pCSII and pCMV-VSVG-RSV-Rev were kindly gifted from H. Miyoshi (RIKEN BioResource Center, Ibaraki, Japan). pGP was gifted from T.Akagi. psPAX2 was the kind gift of Didier Trono (Addgene plasmid: no. 12260). mCherry-Golgi-7 was a gift from Michael Davidson (Addgene plasmid: no. 55052) and the cassette was subcloned into a pCSIIbsr vector. pCAGGS-T2TP was a gift from Koichi Kawakami (National Institute of Genetics, Shizuoka, Japan).The cDNA of mCherry was subcloned into a pCSIIpuro vector to generate pCSIIpuro-mCherry.

### Reagents and antibodies

The following reagents were used: trametinib (no. T-8123, LC Laboratories, Woburn, MA), Y-27632 (no. 253-00513, Wako, Osaka, Japan), PD153035, marimastat (no. SC-202223, Santa Cruz Biotechnology, Dallas, TX), rapamycin (no. R-5000, LC Laboratories), TPA (no. P-1680, LC Laboratories), Rhodamine Phalloidin (no. R415, Invitrogen, Carlsbad, CA, 1:40 dilution for immunofluorescence), EGF (no. E9644, SIGMA, St. Louis, MO).

The following primary and secondary antibodies were used for immunoblotting: Anti-EGFR rabbit antibody (no. 4267, Cell Signaling Technology, Danvers, MA, 1:1,000 dilution); anti-phospho-EGFR (Tyr1068) rabbit antibody (no. 3777, Cell Signaling Technology, 1:1,000 dilution); anti-RAF1 mouse antibody (no. 610152, BD Biosciences, Franklin Lakes, NJ, 1:1,000 dilution); anti-phospho-RAF1 (Ser338) rabbit antibody (no. 9427, Cell Signaling Technology, 1:1,000 dilution); anti-MEK1/2 rabbit antibody (no. 9122, Cell Signaling Technology, 1:1,000 dilution); anti-phospho-MEK1/2 (Ser217/221) rabbit antibody (no. 9121, Cell Signaling Technology, 1:1,000 dilution); anti-ERK1/2 mouse antibody (no. 610123, BD Biosciences, 1:2,000 dilution); anti-phospho-p44/42 MAPK (Erk1/2; Thr202/Tyr204) rabbit antibody (no. 9101, Cell Signaling Technology, 1:2,000 dilution); anti-α-Actin rabbit antibody (no. 4970, Cell Signaling Technology, 1:1,000 dilution); anti-Cdc42 mouse antibody (no. 610929, BD Biosciences, 1:1,000 dilution); anti-Rac1 mouse antibody (no. 610650, BD Biosciences, 1:1,000 dilution); anti-□-catenin mouse antibody (no. 610194, BD Biosciences, 1:1,000 dilution); IRDye 680-conjugated goat anti-mouse IgG antibody (no. 926-32220, LI-COR Biosciences, Lincoln, NE, 1:10,000 dilution); and IRDye 800CW goat anti-rabbit IgG antibody (no. 926-32211, LI-COR Biosciences, 1:10,000 dilution).

The following primary and secondary antibodies were used for immunofluorescence: anti-phospho-Myosin Light Chain 2 (Thr18/Ser19) rabbit antibody (no. 3674, Cell Signaling Technology, 1:50 dilution); anti-E-cadherin rabbit antibody (no. 3195, Cell Signaling Technology, 1:300 dilution); Alexa 647-conjugated goat anti-mouse IgG (H+L) antibody (no. A-21235, Thermo Fisher Scientific, Waltham, MA, 1:1,000 dilution); alexa 647-conjugated goat anti-rabbit IgG (H+L) antibody (no. A-21245, Thermo Fisher Scientific, 1:1,000 dilution); and alexa 568-conjugated goat anti-rabbit IgG (H+L) antibody (no. A-11036, Thermo Fisher Scientific, 1:1,000 dilution).

### Cell culture

MDCK cells were purchased from the RIKEN BioResource Center (no. RCB0995). Lenti-X 293T cells were purchased from Clontech (no. 632180, Mountain View, CA). These cells were maintained in D-MEM (no. 044-29765, Wako, Osaka, Japan) supplemented with 10% FBS (no. 172012-500ML, SIGMA, St. Louis, MO), 100 unit mL^−1^ penicillin, and 100 µg mL^−1^ streptomycin (no. 26253-84, Nacalai Tesque, Kyoto, Japan) in a 5% CO_2_ humidified incubator at 37°C.

### Establishment of stable cell lines

For the establishment of EKAREV-NLS–expressing MDCK cells, a Tol2 transposon system was used. MDCK cells were co-transfected with pT2A-EKAREV-NLS and pCAGGS-T2TP encoding Tol2 transposase, and sorted by FACS as previously described^63,64^. To establish MDCK cells stably expressing 2paRAF, pT2ADWpuro_2paRAF or pT2ADWpuro_2paRAF_ΔmScarlet-I were used for the Tol2 transposon system. For the generation of cells stably expressing Eevee-ROCK-NES and the other ectopic proteins, a lentiviral or retroviral system was employed. To prepare the lentivirus, pCSII-based lentiviral vector^65^ or lentiCRISPRv2 (Addgene Plasmid: no. 52961), psPAX2 (Addgene Plasmid: no. 12260), and pCMV-VSV-G-RSV-Rev were co-transfected into Lenti-X 293T cells using polyethylenimine (no. 24765-1, Polyscience Inc., Warrington, PA). To prepare the retrovirus, pCX4-based retroviral vector^66^ or pSUPER (Oligoengine, Seattle, WA), pGP, and pCMV-VSV-G-RSV-Rev were co-transfected into Lenti-X 293T cells. Stable cell lines of MDCK cells were selected and maintained in media containing the following antibiotics: MDCK/EKAREV-NLS/2paRAF, 4 µg mL^−1^ puromycin (no. P-8833, SIGMA); MDCK/EKAREV-NLS/2paRAF/Golgi-7-mCherry, 4 µg mL^−1^ puromycin and 10 µg mL^−1^ blasticidin S (no. 029-18701, Wako); MDCK/Eevee-ROCK-NES, 10 µg mL^−1^ blasticidin S; MDCK/Eevee-ROCK-NES/2paRAF, 10 µg mL^−1^ blasticidin S and 4 µg mL^−1^ puromycin; MDCK/EKAREV-NLS/mCherry, 4 µg mL^−1^ puromycin; MDCK/RA Rho GEF, 4 µg mL^−1^ puromycin and 10 µg mL^−1^ blasticidin S.

### CRISPR/Cas9-mediated KO cell lines

For CRISPR/Cas9-mediated KO of dog *CTNNA1* (α-catenin), single guide RNAs (sgRNA) targeting the exon were designed using the CRISPRdirect^67^. The following sequence was used for the sgRNA sequence: GTAGAAGATGTTCGAAAACA. Oligo DNAs for the sgRNA were cloned into the lentiCRISPRv2 vector, and the sgRNA and Cas9 were introduced into MDCK cells by lentiviral infection. The infected cells were selected with 4.0 µg mL^−1^ puromycin. After the selection, reduction in expression levels of the proteins was confirmed by immunoblotting. Bulk cells were used for the experiments.

### shRNA-mediated KD cell line

For shRNA-mediated KD of dog *RAC1* and *CDC42*, the DNA oligomers corresponding to the shRNAs targeting the genes were subcloned into pSUPER vector. The following sequences were used for shRNA target sequences: *RAC1*, GCCTTCGCACTCAATGCCAAG; *CDC42*, GAACAAACAGAAGCCTATC. The shRNAs were introduced into MDCK cells by retroviral infection. The infected cells were selected with 4.0 µg mL^−1^ puromycin. After the selection, reduction in expression levels of the target proteins was confirmed by immunoblotting.

### Confinement release assay

To confine the MDCK cell monolayer, a Culture-Insert 2 Well (no. 81176, ibidi, Martinsried, Germany) was placed on a glass-bottom dish coated with 0.3 mg mL^−1^ type I collagen (Nitta Gelatin, Osaka, Japan). MDCK cells (7 × 10^3^ cell) were seeded in the Culture-Insert. 24 h after seeding, the Culture-Insert was removed, and the medium was replaced with Medium 199 (11043023; Life Technologies, Carlsbad, CA) supplemented with 10% FBS, 100 unit mL^−1^ penicillin, and 100 µg mL^−1^ streptomycin. 30 min after the removal of the Culture-Insert, the cells were imaged with an epifluorescence microscope every 1 to 10 min.

### Boundary assay

For the light-induced ERK activation experiment, MDCK/EKAREV-NLS/2paRAF cells were seeded in a well of a Culture-Insert 2 well placed on a 24 well glass bottom plate, and MDCK/EKAREV-NLS cells were seeded in the other well of the insert and outside of the insert. After 15 h incubation, the insert was removed, followed by further incubation for 36 h, allowing the cells to fill the gap between the cell populations. The interface between cells with and without 2paRAF expression was imaged, and the cells were exposed to 438 nm blue LED light every 5 min for CRY2 activation.

For the drug-induced cell contraction experiment, a well of the insert was removed by cutting, and the remained insert with a well was placed on a 24 well glass-bottom plate. MDCK cells expressing RA Rho GEF were seeded in the well. After 6 h incubation, the insert was removed, and MDCK/EKAREV-NLS cells were plated in the 24 well glass-bottom plate. The interface between cells with and without RA Rho GEF expression was imaged, and dimethyl sulfoxide (DMSO; final 0.1%) or rapamycin (final 50 nM) was added into the medium for the Rho GEF activation. To determine the ERK wave directionality, heat maps of ERK activity were obtained by interpolating the signals in regions between the nuclei of MDCK/EKAREV-NLS cells in the FRET/CFP ratio images. The heat maps of ERK activity were analyzed by particle image velocimetry (PIV) using a free Matlab-toolbox, MatPIV (a GNU public license software, https://www.mn.uio.no/math/english/people/aca/jks/matpiv/), to calculate the ERK wave directionality. The size and overlap of the interrogation window was set to 349 µm and 75%, respectively. To determine the directionality of cell displacement, the Fiji TrackMate plugin^68,69^ was applied to the CFP fluorescence images for tracking each cell over 13-13.5 h after treatment with DMSO or rapamycin.

### Cell stretch assay

For the cell stretch assay, MDCK cells (2 × 10^5^ cell) were seeded on an elastic silicone chamber (no. STB-CH-04, STREX, Osaka, Japan) coated with 0.3 mg mL^−1^ type I collagen. After 24 h incubation, the MDCK cells on the stretch chamber were uniaxially stretched by 50% with a manual cell-stretching system (no. STB-100-04, STREX) on an epifluorescence microscope. For immunoblotting, stretched MDCK cells were lysed with SDS sample buffer containing 62.5 mM Tris-HCl (pH6.8), 12% glycerol, 2% SDS, 40 ng mL^-1^ bromophenol blue, and 5% 2-mercaptoethanol, followed by sonication with a Bioruptor UCD-200 (Cosmo Bio, Tokyo, Japan). After boiling at 95°C for 5 min, the samples were resolved by SDS-PAGE on SuperSep Ace 5-20% precast gels (Wako), and transferred to PVDF membranes (Merck Millipore, Billerica, MA). All antibodies were diluted in Odyssey blocking buffer (LI-COR Biosciences). Proteins were detected by an Odyssey Infrared Imaging System (LI-COR Biosciences).

### Time-lapse imaging

FRET images were obtained and processed under essentially the same conditions and procedures as previously described^70^. Briefly, cells were imaged with an IX83 inverted microscope (Olympus, Tokyo, Japan) equipped with a UPlanFL-PH 10x/0.3 (Olympus), a UPlanSApo 20x/0.75 (Olympus), or a UPlanSApo 40x/0.95 objective lens (Olympus), a DOC CAM-HR CCD camera (Molecular Devices, Sunnyvale, CA), a Spectra-X light engine (Lumencor Inc., Beaverton, OR), an IX3-ZDC laser-based autofocusing system (Olympus), an electric XY stage (Sigma Koki, Tokyo, Japan), and a stage top incubator (Tokai Hit, Fujinomiya, Japan). The filters and dichromatic mirrors used for time-lapse imaging were as follows: for FRET imaging, a 438/24 excitation filter (incorporated in the Spectra-X light engine), a FF458-Di02-25×36 (Semrock, Rochester, NY) dichromatic mirror, and FF01-483/32-25 (Semrock) and FF01-542/27-25 (Semrock) emission filter for CFP and FRET, respectively. For mCherry and mScarlet-I imaging, a 575/25 excitation filter, a glass dichromatic mirror (Olympus), and FF01-624/40-25 (Semrock) emission filters were used.

### Immunofluorescence and confocal microscopy

MDCK cells were fixed with 4% paraformaldehyde in PBS for 15 min at room temperature, followed by permeabilization with 0.2% Triton X-100 in PBS for 5 min. The samples were then incubated with 1% BSA in PBS for 1 h at room temperature, followed by sequential incubation with primary and secondary antibodies diluted with 1% BSA in PBS overnight at 4°C (primary antibodies) or for 1 h at room temperature (secondary antibodies). Images were collected using a Fluoview FV1000 confocal microscope (Olympus) equipped with a UPlanSApo 60x/1.35 or a UPlanSApo 100x/1.40 objective lens (Olympus).

### Traction force microscopy

Polyacrylamide gel substrates were prepared as previously described, with slight modifications ^24,28^. Briefly, glass-bottom dishes (IWAKI, Shizuoka, Japan) were treated with 2% acetic acid (WAKO) and 0.2% 3-(Trimethoxysilyl)propyl methacrylate (SIGMA) in 80% ethanol for 2 min. After the removal of the solution, the dishes were dried for 15 min. For 3 kPa gels, 5.5% acrylamide, 0.09% bisacrylamide, 0.05% ammonium persulfate, 0.05% N,N,N’,N’-tetramethyl ethylenediamine, and 0.01% deep red fluorescent carboxylate-modified beads (0.2 µm diameter; F8810, Thermo Fisher Scientific) in PBS were prepared. 18 µL of the solution was put on the glass-bottom dishes and 18 mm glass cover slips (Matsunami, Osaka, Japan) were placed on top of them. After polymerization, the gels were covered with 2 mg mL^-1^ Sulfo-SANPAH (no. ab145610, abcam, Cambridge, UK) and activated by ultraviolet light for 5 min with an Ultraviolet Crosslinker CL-1000 apparatus (UVP, Upland, CA). The procedure was repeated again with 7 min UV irradiation. The gels were coated with 100 µm mL^-1^ type I collagen overnight at 4°C. Then, the gels were washed three times with PBS and incubated with culture medium for 1h. For traction force microscopy, the bead fluorescence was imaged with an IX83 inverted microscope equipped with a 632/22 excitation filter (incorporated in the Spectra-X light engine), a glass dichromatic mirror (Olympus), and FF01-692/40-25 (Semrock) emission filter. A reference image was obtained after the removal of cells by trypsinization. Traction forces were computed by Fourier-transform traction microscopy as described previously^13^.

### Quantification of ERK activity and cell deformation

For the analysis of the cell strain rate along an axis of collective cell migration (x-strain rate), PIV using MatPIV was applied to phase contrast time-lapse images to calculate velocity fields of cells. Velocity fields at time *T* were computed by the displacement between *T*-*Δt* and *T+Δt*. The size of the interrogation window was set to 29.1 µm, approximately corresponding to the typical cell size, and the window overlap was set to 50%. Then, the x-strain rate was calculated according to

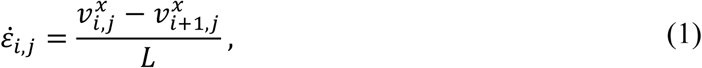

where 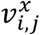 is the *x* component of the velocity at spatial indices of resultant windows obtained by PIV analysis (*i, j*), and *L* is the distance between the windows (14.5 µm).

To represent the FRET efficiency, FRET/CFP ratio images were generated after the background intensity was subtracted from the original fluorescence images in the CFP and FRET channel, respectively, using Metamorph software (Molecular Devices, Sunnyvale, CA). Then, the Fiji TrackMate plugin was applied to the CFP fluorescence images for tracking each cell. The time derivative of ERK activity (ERK activation rate) at time *t* for each cell was computed by the change in FRET/CFP ratio between *t*-*Δt* and *t+Δt*. The obtained time-series data of the x-strain rate and the ERK activation rate were processed with a Savitzky-Golay filter to reduce the noise. The cross-correlation coefficient *r*(*τ*) between time-series data of x-strain rate *f*(*t*) and ERK activation rate *g*(*t*) was calculated as follows:

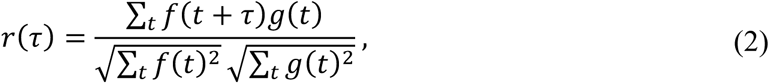

where *τ* is the lag time.

To quantify the amplitude of x-strain rate oscillation, the time-series data were fitted to a multi-peak function as described previously^71^.

### Quantification of directedness

For quantification of directionality of Golgi positioning, ERK activation waves, and cell migration, we defined directedness as shown in Fig. 6f. The reference direction is set to the forward direction for analysis of Golgi orientation and cell migration direction, and to the backward direction for that of ERK wave direction to match the signs of their directedness. To obtain the sample directions of the Golgi apparatus, the center of mass of the nucleus in each cell was determined with Fiji. Fluorescence images of Golgi apparatus markers were processed with a mean filter. The position that showed the highest intensity in the region within 6.8 µm from the center of mass of the nucleus was defined as the position of the Golgi apparatus. The sample directions were determined by the direction of the Golgi apparatus relative to the center of mass of the nucleus. To determine the sample directions of ERK activation waves, heat maps of ERK activity were obtained by interpolating the signals in regions between the nuclei of MDCK/EKAREV-NLS cells in the FRET/CFP ratio images. The heat maps of ERK activity were analyzed by MatPIV with a 174 µm window size and a 75% window overlap. The directions of the calculated velocity vectors were obtained as the sample directions. For the determination of the sample directions of cell migration, each cell was tracked with a Fiji TrackMate plugin. The direction of cell displacement for 20 min was defined as the sample direction of cell migration for that individual cell.

### Optogenetic ERK activation wave assay

To illuminate a defined rectangular region with blue light, transillumination light of a 100-W halogen lamp filtered with BA420-460 (Olympus) was covered by a homemade aperture mask containing a slit. The image of the slit was focused on the sample plane by the condenser lens. The width of the focused image of the illumination light was approximately 35 µm. The illuminated region was moved at 3 µm min^-1^ velocity, and the illumination was patterned with 400 µm spatial intervals at each time point. The procedure was applied to the migrating MDCK/EKAREV-NLS/2paRAF/Golgi-7-mCherry cells treated with 1 µM PD153035, an EGFR inhibitor, to suppress autonomous ERK activation waves.

### Kymography

To obtain the kymographs of FRET/CFP ratios and x-strain rate, these values were averaged along the y-axis in a defined region of the images, providing an intensity line along the x-axis. The operation was repeated for the respective time points, and the intensity lines were stacked along the y-axis for all time points.

### Statistical analysis

Statistical analyses were performed with GraphPad Prism 7 software (GraphPad Software, San Diego, CA). No statistical analysis was used to predetermine the sample size. The sample sizes, statistical tests, and p-values are indicated in the figures and the figure legends. To compare two sets of data, paired or unpaired *t*-tests were used. P-values of less than 0.05 were considered to be statistically significant in two-tailed tests, and were classified as follows: *P < 0.05, **P < 0.01, ***P < 0.001, ****P < 0.0001, and n.s. (not significant, i.e., P ≥0.05).

### Mathematical model and simulation

See the supplementary note for details.

## Supporting information

mov1

mov2

mov3

mov4

mov5

mov6

mov7

Mov8

Mov9

Mov10

## Code availability

Computer codes developed for this study are available on request to the corresponding authors.

## Repeatability of experiments

All experiments were performed on at least three independent cell culture preparations.

## Acknowledgements

This work was supported by JSPS KAKENHI Grant Numbers 17J02107, 15H05949 “Resonance Bio”, by the SPIRITS 2018 of Kyoto University, by Inamori Research Grants, and by the Kyoto University Live Imaging Center. We would like to thank Manuel Gómez González regarding the analysis of traction force microscopy, James-Alan Hejna for English editing, and Daniel Boocock, Edouard Hannezo and Naoki Honda for fruitful discussions.

## Author contributions

Conceptualization, N.H., M.M., T.H.; Methodology, N.H., K.A., X.T., M.M., T.H.; Software, N.H., X.T., T.H.; Validation, N.H., L.R., A.M.L., T.H.; Formal analysis, N.H., T.H.; Investigation, N.H., L.R., A.M.L.; Resources, N.H., K.A., X.T., T.H.; Data curation, N.H., T.H.; Writing - original draft, N.H., M.M., T.H.; Writing - review & editing, N.H., K.A., M.M., T.H.; Visualization, N.H., T.H.; Supervision, M.M., T.H.; Project administration, M.M., T.H.; Funding acquisition, N.H., X.T., M.M., T.H.

## Materials & Correspondence

should be addressed to the corresponding authors.

## Supplementary Information

**Extended Data Fig. 1: EGFR and ADAM activity are required for ERK activity propagation**

**a**-**c**, Kymographs of ERK activity during collective cell migration in MDCK cells treated with DMSO (**a**), 1 µM PD153035 (**b**), or 10 µM marimastat (**c**). The inhibitors were added at 3.5 h after the start of time-lapse imaging.

**Extended Data Fig. 2: ERK activation is required for cell deformation during collective cell migration**

**a**-**c**, x-strain rate of migrating cells treated with DMSO (**a**), 1 µM PD153035 (**b**), or 200 nM trametinib (**c**) are plotted. Each color indicates three representative cells. **d**, Mean amplitude of the x-strain rate oscillation in each cell treated with DMSO, PD153035, or trametinib is plotted. Red lines indicate the mean of the mean amplitude. *n* = 326 cells (DMSO), 445 cells (PD153035), and 275 cells (trametinib) from three independent experiments. Unpaired *t*-test, *P* < 0.0001.

**Extended Data Fig. 3: Traction force generation gradually increases as time elapses without optogenetic ERK activation**

**a**, Traction force microscopy of cells with and without 2paRAF expression. Representative DIC images (upper), CIBN-mScarlet-I-CAAX (2paRAF; middle), and traction forces (lower) are shown at 0 h (left), 8 h (center), and 12 h (right) after the start of time-lapse imaging. **b**, Quantification of mean traction forces under the cells with and without 2paRAF expression in (**a**), with SDs (n = 3).

**Extended Data Fig. 4: Migrating cells possess front-rear polarity toward the direction of migration**

**a**, Immunofluorescence images of F-actin (left) and di-phosphorylated MLC (ppMLC; center) at the apical plane in MDCK cells at 17 h after migration. At the right, the images of F-actin and ppMLC are merged. **b**, Immunofluorescence images of F-actin (left) and E-cadherin (center) at the apical (upper row) and basal (lower row) planes in MDCK cells at 17 h after migration. **c**, Immunofluorescence images of the Golgi apparatus (GM130) and nucleus (EKAREV-NLS) in MDCK cells immediately after removing confinement (0 h migration). The upper image shows a wide-view field, and the lower images are magnified regions corresponding to the numbered windows in the upper image. **d**, Polar histograms showing the distribution of the Golgi orientation relative to the nucleus in cells at 0-0.5 mm (left), 0.5-1.0 (center), and greater than 1.0 mm (right) distant from the leader cells. *n* = 1575 cells (0-0.5 mm), 1529 cells (0.5-1.0 mm), and 958 cells (>1.0 mm) from three independent experiments. **e**, Analysis of the expression level of Cdc42 and α-actin in WT and shCdc42-expressing MDCK cells by immunoblotting. **f**, Analysis of the expression level of Rac1 and α-actin in WT and shRac1-expressing MDCK cells by immunoblotting.

**Extended Data Fig. 5: ERK activation waves are required for multicellular alignment of front-rear cell polarity**

**a**, Immunofluorescence images of the Golgi apparatus (GM130) and nucleus (EKAREV-NLS) in α-catenin KO MDCK cells after 21 h migration. An upper image shows a wide-view field, and the lower images are magnified images of the region indicated in the upper image with numbered white rectangles. **b**, Polar histograms showing the distribution of Golgi orientation relative to the nucleus in the cells at 0-0.5 mm (left), 0.5-1.0 mm (center), and greater than 1.0 mm (right) distant from the leader cells. *n* = 572 cells (0-0.5 mm), 1564 cells (0.5-1.0 mm), and 957 cells (>1.0 mm) from three independent experiments. **c**, Mean directedness of Golgi orientation binned every 300 µm from the leader cells in (**b**) is plotted as a function of distance from the leader cells, with SDs. The data were obtained from three independent experiments. **d, e**, Kymographs of ERK activity during collective cell migration in MDCK cells treated with 200 nM trametinib (**d**) and 10 nM TPA (**e**). These inhibitors were added 3.5 h after the start of time-lapse imaging.

**Extended Data Fig. 6: Proposed model underlying long-distance transmission of directional cues for collective cell migration**

(A) Each migrating cell possesses an ERK-mediated mechanochemical feedback system. Extension of a cell activates ERK, and the activated ERK trigger cell contraction, forming a negative feedback loop between cell extension and ERK activation. (B) Advancement of the leader cells toward the free space exerts pulling forces onto the adjacent follower cells, directing the front-rear cell polarization (i). The pulling force then stretches the follower cells and activates ERK in the follower cells (ii). The activated ERK triggers polarized rear contraction, evoking another round of cell extension, front-rear polarization, and ERK activation in the next set of follower cells (iii). Thus, the chain propagation of ERK activation and contractile force generation spread over a long distance, leading to multicellular alignment of front-rear cell polarity, i.e., collective cell polarization (iv). Illustrated by Hiroko Uchida.

**Supplementary Video 1: ERK activation waves and cell deformation waves during collective cell migration**

Time-lapse video of collectively migrating MDCK cells expressing EKAREV-NLS. Phase contrast images are shown at the top. The golden pseudo-color represents the FRET/CFP ratio indicating ERK activity (middle). Red and blue indicate positive and negative x-strain rate, respectively (bottom). The color scales correspond to those in Fig. 1a. Time in hr:min.

**Supplementary Video 2: Cell contraction upon optogenetic ERK activation**

Time-lapse video of the boundary between confluent MDCK cells with and without 2paRAF expression. The upper frames represent the fluorescence of CIBN-mScarlet-I-CAAX, a component of 2paRAF. The lower images show differential interference contrast (DIC). The blue light illumination started at 0 min and was repeated every 5 min. The cells were treated with DMSO (left), 200 nM trametinib (center), 10 µM Y-27632 (right) 30 min before the start of the imaging. Time in hr:min.

**Supplementary Video 3: ROCK activity propagation during collective cell migration**

First part: Time-lapse video of collectively migrating MDCK cells expressing a FRET biosensor for ROCK activity. The color represents the FRET/CFP ratio indicating ROCK activity and its scale corresponds to the one in Fig. 3f. Second part shows migrating cells that were treated with 200 nM trametinib (left) and 100 µM Y-27632 (right) at 0 min. Time in hr:min.

**Supplementary Video 4: Traction force microscopy with optogenetic ERK activation** Time-lapse traction force microscopy at the boundary between confluent MDCK cells with and without 2paRAF expression. Upper frames show differential interference contrast (DIC). Middle frames represent the fluorescence of CIBN-mScarlet-I-CAAX, a component of 2paRAF. Lower frames show traction forces. The color scale corresponds to the one in Fig. 4a. The blue light illumination started at 0 min and was repeated every 5 min. Time in hr:min.

**Supplementary Video 5: Cell contraction triggers ERK activation waves**

Time-lapse videos of the boundary between confluent MDCK cells with and without rapamycin-activatable Rho GEF (RA Rho GEF). The color represents the FRET/CFP ratio indicating ERK activity, and the scale corresponds to the one in Fig. 5b. The cells were treated with DMSO (left) or 50 µM rapamycin (right) at 0 min to induce contraction of the RA Rho GEF-expressing cells. Time in hr:min.

**Supplementary Video 6: Intercellular mechanical linkage is required for ERK activation waves**

Time-lapse videos of collectively migrating WT (left) and α-catenin KO (right) MDCK cells. The color represents the FRET/CFP ratio indicating ERK activity, as shown in Fig. 5, and its upper and lower value is 1.8 and 0.85. Time in hr:min.

**Supplementary Video 7: Requirement of Cdc42 and Rac1 for unidirectional ERK activation waves**

Time-lapse videos of collectively migrating WT (upper), shCdc42 (middle), and shRac1 (lower)-expressing MDCK cells. The color represents the FRET/CFP ratio indicating ERK activity, as shown in Fig. 1, and its upper and lower value is 1.8 and 0.85 for each of the videos. Time in hr:min.

**Supplementary Video 8: Effect of constitutive ERK activation and inhibition on collective cell migration**

Time-lapse videos of collectively migrating MDCK cells. The color represents the FRET/CFP ratio indicating ERK activity, and the scale corresponds to the one in Extended Data Fig. 6d,e. The cells were treated with DMSO (upper), 200 nM trametinib (middle), and 10 nM TPA (bottom) at 0 min. Time in hr:min.

**Supplementary Video 9: Optogenetic ERK activation waves and front-rear cell polarization**

Time-lapse videos of migrating MDCK cells without (upper) and with (lower) Optogenetic ERK activation waves. Green and red indicate nuclei (EKAREV-NLS) and Golgi apparatus (Golgi-7-mCherry), respectively. Cyan represents regions illuminated with blue light. The cells were pre-treated with 1 µM PD153035, an EGFR inhibitor, to suppress autonomous ERK activation waves. Time in hr:min.

**Supplementary Video 10: in silico ERK activation waves**

Simulations for ERK activation waves in migrating cells with (ω=1, upper frame) and without (ω=0, lower frame) cell polarity. The color represents the ERK activity and its scale corresponds to the one in Fig. 8c.

