## supplementary_figure for "ERK-mediated mechanochemical waves direct collective cell polarization"

Extended Data Fig. 1

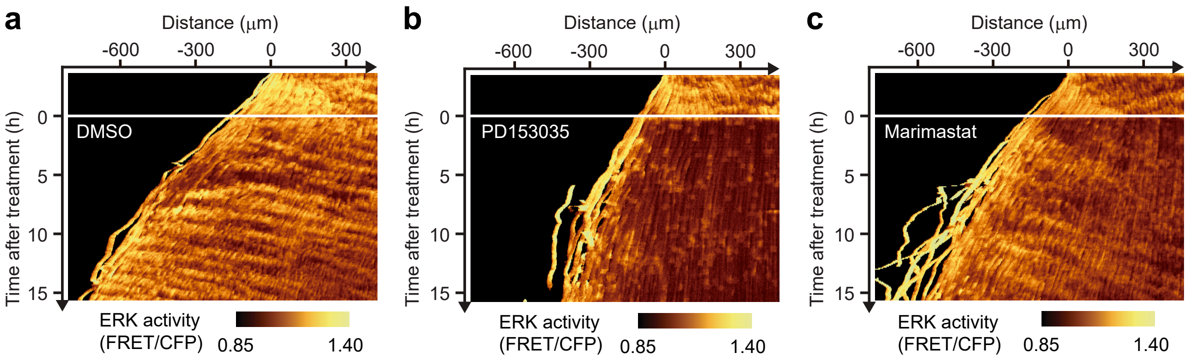


Hino et al.

Extended Data Fig. 2

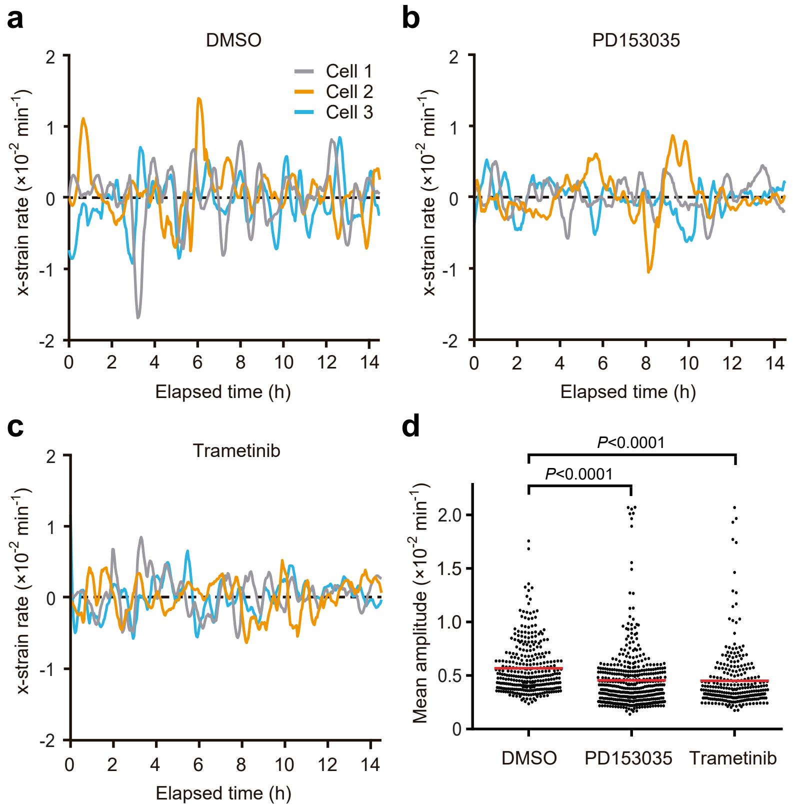


Hino et al.

Extended Data Fig. 3

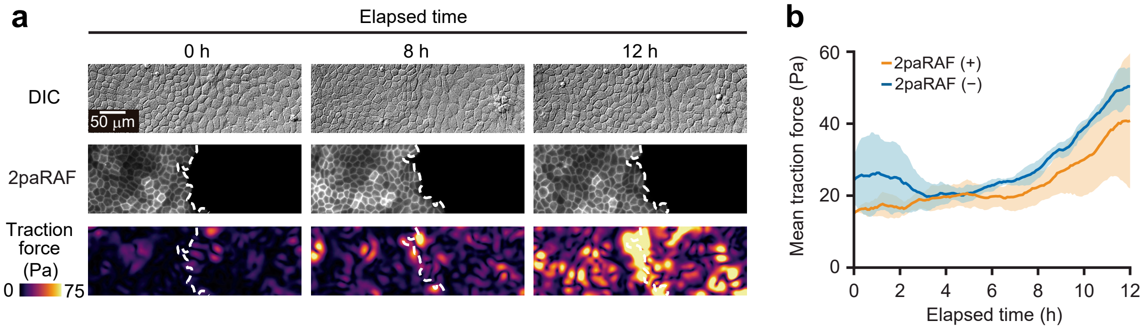


Hino et al.

Extended Data Fig. 4

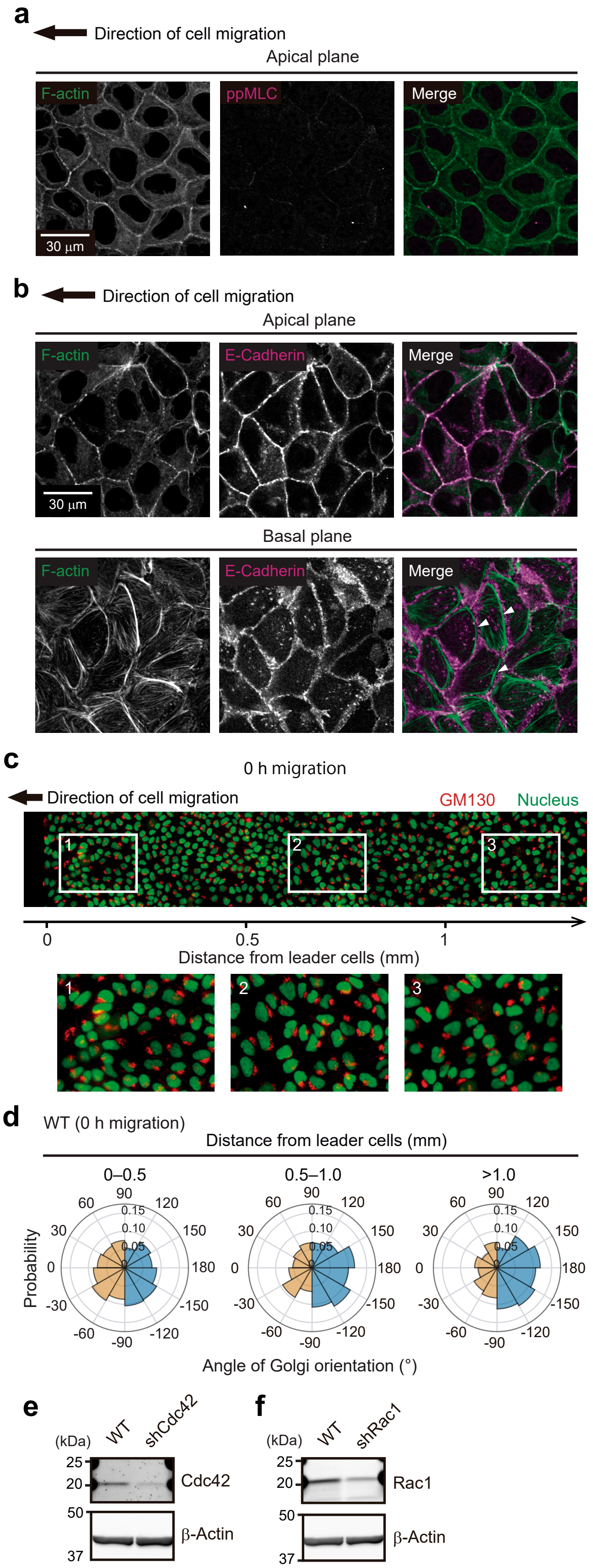


Hino et al.

### Extended Data Fig. 5

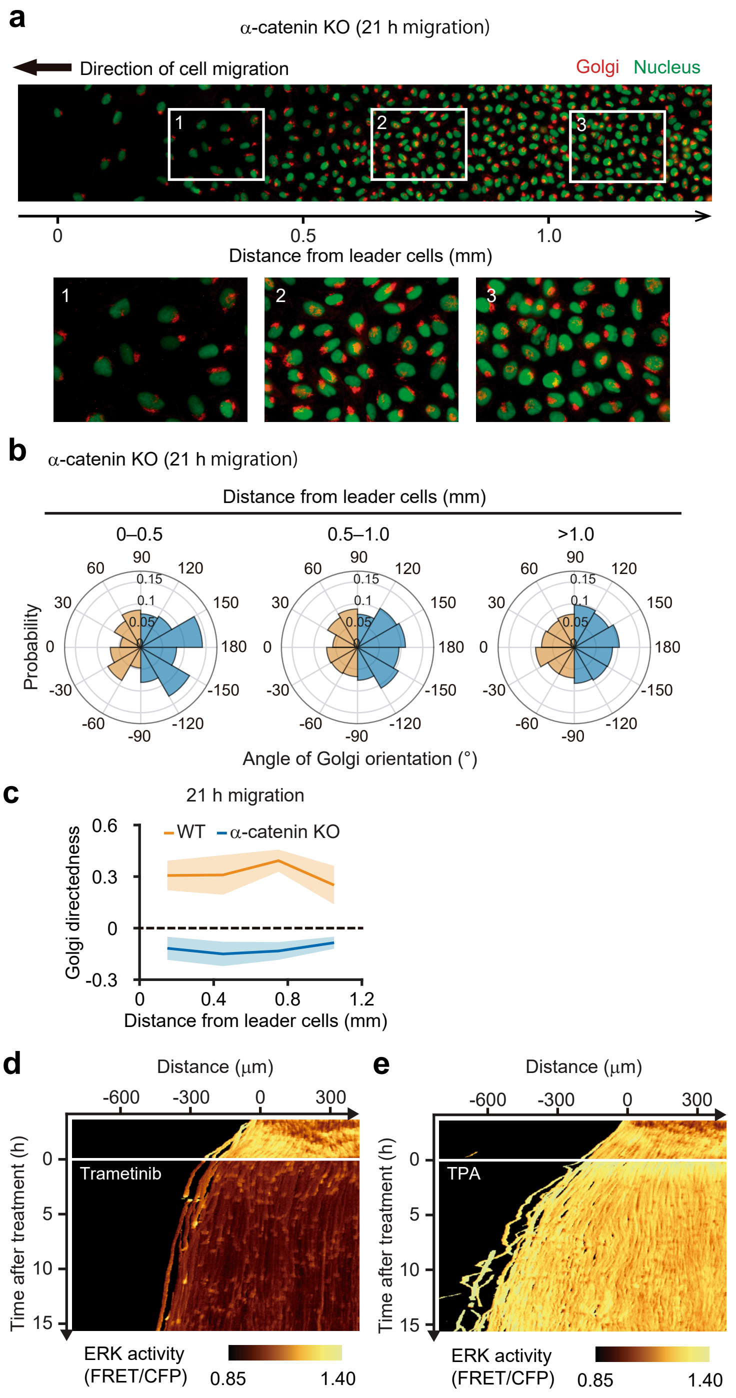


Hino et al.

Extended Data Fig. 6

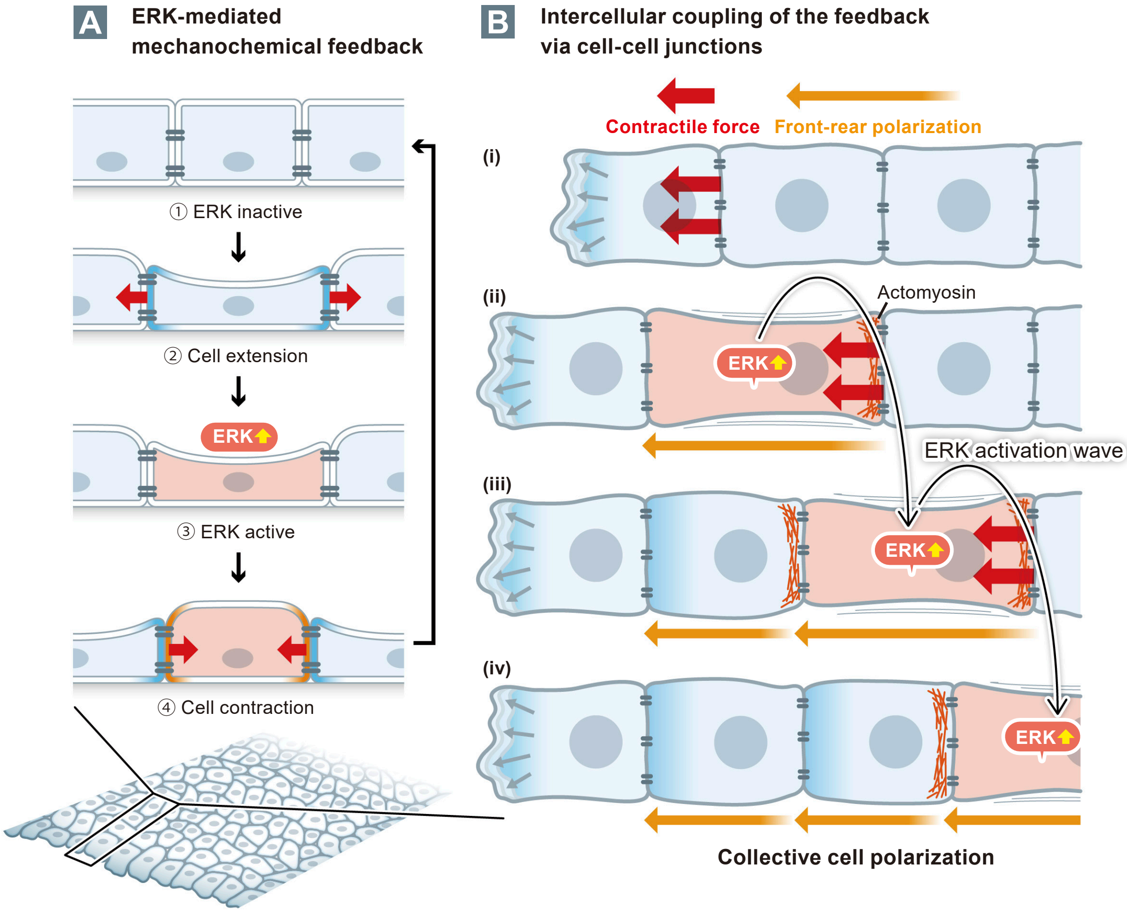
