## supplementary note for "ERK-mediated mechanochemical waves direct collective cell polarization"

### Supplementary notes on a mathematical model and numerical analysis

Hino, N., et al.

#### 1. A cellular Potts model for cell migration

We used a two-dimensional cellular Potts model (CPM), also known as a Glazier-Graner-Hogeweg model, widely used for multicellular dynamics (Balter et al., 2007; Krieg et al., 2008; te Boekhorst et al., 2016; van Helvert et al., 2018). In the CPM, each cell morphology is represented as a cluster of square lattices identified with the identical index $\sigma$, denoting a cell; the lattice distribution mainly determines an energy of the system *H*, such that cell behavior depends on a balance of forces defined by the energy. The energy in our model is composed of minimal factors necessary to capture the two-dimensional multicellular dynamics, such as interfacial energy, cell area constraint, and active cell contraction as follows:

$$H = \sum_{\boldsymbol{r},\boldsymbol{r}'} J_{\tau(\sigma_{\boldsymbol{r}})\tau(\sigma_{\boldsymbol{r}'})}\left(1 - \delta_{\sigma_{\boldsymbol{r}}\sigma_{\boldsymbol{r}'}}\right) + \lambda_A \sum_{\sigma} (A_\sigma - A_0)^2 + H_{contraction}, \tag{1}$$

where each of $\boldsymbol{r}$ and $\boldsymbol{r}'$ represents a position of lattice site, $\tau$ is an attribute of the lattice, i.e., cell or medium, *J* is the interfacial energy between cell-cell or cell-medium, $\delta$ is the Kronecker delta, $\lambda_A$ is the magnitude of resistance to cell deformation, $A_\sigma$ is the current cell area, $A_0$ is the ideal cell area, and $H_{contraction}$ is a term for cell contraction.

The first term in Eq. (1) describes a strength of cell-cell adhesion. The energy to maintain cell-cell adhesion is determined by an energy of cell-cell adhesion $J_{cc}$ relative to that of cell-medium adhesion $J_{cm}$ and is expressed as $\gamma = J_{cm} - J_{cc}/2$ (Davies and Rideal, 1963; Glazier and Graner, 1993). We determined $J_{cm} = J_{cc} = 3$ in light of a balance with other parameters for imitating MDCK behaviors, and changed the value in $J_{cm}$ for the change in $\gamma$. The second term represents cell elasticity, meaning that the cells attempt to retain the ideal area. We set $\lambda_A$ as 0.02 and 1 when cells extend beyond and shrink below the ideal area, respectively, because of the unique material property of cells (Latorre et al., 2018; Trepat and Sahai, 2018). The ideal cell area was determined as $A_0$= 400 μm$^2$, by the use of actual cell images. The third term relates to the active cell contraction. In this model, individual cells are assigned unit vectors $\boldsymbol{p_\sigma}$ corresponding to the front-rear cell polarity, and the cell contraction occurs depending on the polarity level $\omega$ with the contraction strength $\lambda_\sigma$, a variable related to the ERK activity as shall be explained later. We define the contraction term as follows:

$$H_{contraction} = \sum_{\sigma} \sum_{l_\sigma} \lambda_\sigma \Phi, \tag{2}$$

and

$$\Phi = \begin{cases} 1 + 2\omega\pi^{-1}(\varphi_{l_\sigma} - \pi) & \text{if} \quad \varphi_{l_\sigma} > \pi(1 - (2\omega)^{-1}) \\ 0 & \text{if} \quad \varphi_{l_\sigma} \leq \pi(1 - (2\omega)^{-1}) \end{cases}, \tag{3}$$

where $l_\sigma$ represents the index of lattices composing the cell periphery, and $\varphi_{l_\sigma} \in [0, \pi]$ is an angle between $\boldsymbol{p_\sigma}$ and the vector connecting from the centroid of a cell to a peripheral lattice. Note that $H_{contraction} = \sum_\sigma \lambda_\sigma P_\sigma$, where $P_\sigma$ is the perimeter of a cell when $\omega = 0$, meaning that cells shrink independent of the direction of polarity. By contrast, cell contraction is biased to the rear of the cell when $\omega$ becomes larger.

In the CPM, the system transition occurs stochastically by a lattice-based Monte Carlo method; that is, the labeled value of a randomly chosen lattice site $\sigma_{\boldsymbol{r}}$ is attempted to be replaced by a different labeled value of randomly-chosen adjacent lattice site $\sigma_{r'}$. The transition occurs by evaluating a change in energy $\Delta H$ associated with its replacement. In the case of energy increase, i.e., $\Delta H$>0, the index replacement occurs stochastically according to a Boltzmann acceptance function $exp(-\Delta H)$, while it deterministically occurs in the case of energy decrease, i.e., $\Delta H$≤0. For the details of CPM, see (Glazier and Graner, 1993; Graner and Glazier, 1992; Hirashima et al., 2017; Marée et al., 2007; Merks and Glazier, 2005; Scianna, 2015). We regard the number of trials for lattice replacement as a total number of pixel domains in simulations as a unit simulation step (USS); an update at each lattice site is attempted once per 1 USS on average. We then defined 100 USS as 1 Monte Carlo step, corresponding to 1 hour.

#### 2. Dynamics of polarity orientation

Explicit rules that govern the dynamics of front-rear cell polarity have not been clear. Yet, earlier experimental studies have proposed that the front-rear polarity is oriented according to the tensile force on cell-cell junctions (Das et al., 2015; Hayer et al., 2016). In particular, it has been shown that Merlin localized at a cell-cell junction inhibits the activation of Rac1 when the tension is low. In contrast, with high tension by strong contraction of neighbor cells, Merlin is released into the cytoplasm, and Rac1 is locally activated (Das et al., 2015). Thus, orientation of the front-rear polarity in individual cells changes through physical interaction with neighboring cells over time. With this fact, we model the dynamics of cell polarity orientation $\vartheta_\sigma$ as a phenomenological coupling with cell displacement according to earlier theoretical studies (Hirashima et al., 2013; Notbohm et al., 2016; Peyret et al., 2019; Szabó et al., 2010a; Szabó et al., 2010b; Tlili et al., 2018):

$$\frac{d\vartheta_\sigma}{dt} = \mu \frac{v_\sigma}{\sqrt{A_\sigma}} \Delta\phi_\sigma, \tag{4}$$

where $\mu$ is the degree of alignment to polarity in neighbor cells, $v_\sigma$ is the cell speed, and $\Delta\phi_\sigma$ is the angle between cell velocity $\boldsymbol{v}_\sigma$ and the polarity vector $\boldsymbol{p}_{\boldsymbol{\sigma}}$. $v_\sigma/\sqrt{A_\sigma}$ contributes to weighting of how much the neighboring cells affect reorientation of cell polarity and/or self-reinforcement for a persistent polarization.

The behavior of cell polarity according to Eq. (4) can vary. For example, consider a situation in which a cell A pulls an adjacent cell B in a direction opposite to the polarity within cell B, and the center position of cell B is moved toward cell A. When $\mu$ is larger, the polarity in cell B will re-orient to the direction in which cell B is pulled. Conversely, when $\mu$ is smaller, the polarity orientation in cell B will not change, and it tends to be persistent over time. In addition to values of $\mu$, the magnitude of the cell displacement resulting from interactions between multiple neighboring cells affects multicellular alignment of polarity. We chose $\mu = 1$ for reproducing a proper polarity alignment.

#### 3. ERK-mediated mechanochemical feedback

In the text, we have shown experimentally that cell extension activates ERK, and that the ERK activation induces cell contraction. Here we incorporated this observation into the framework of CPM.

We define the cell areal strain as $\varepsilon_\sigma = A_\sigma/A_0 - 1$, and the dynamics of a normalized ERK activity level $[ERK]_\sigma$, bounded from -1 to 1, is represented as

$$\frac{d[ERK]_\sigma}{dt} = (\tanh(\alpha\varepsilon_\sigma) - [ERK]_\sigma)/\eta_E, \quad (5)$$

where $\alpha$ is a sensitivity parameter and $\eta_E$ denotes a timescale of this dynamics. We set $\alpha = 3$ for a proper response to cell areal strains in simulations, and $\eta_E = 3$ min with measured data in Figure 1.

The dynamics of cell contraction strength $\lambda_\sigma$ is defined as

$$\frac{d\lambda_\sigma}{dt} = (\lambda U([ERK]_\sigma - [ERK]^*) - \lambda_\sigma)/\eta_c, \quad (6)$$

where $\lambda$ is a controlling parameter for conversion of ERK activation to contraction, $[ERK]^*$ denotes a threshold of ERK activation-induced contraction, and $\eta_E$ denotes a timescale of this dynamics. $U(x)$ is a step function: $U(x) = 1$ for $x \geq 0$ and $U(x) = 0$ for $x < 0$. We chose $\lambda = 10$ and $[ERK]^* = 0.5$ since the ERK activation waves are reproduced with them. We set $\eta_c = 30$ min, reflecting the response time delay observed in Figure 3C.

#### 4. Simulations

##### *4-1. Matching time*

All dynamics on polarity orientation, ERK activity, and contraction (Eq. (4)-(6)) were calculated in their discretized form and were updated every USS. The cell velocity $\boldsymbol{v}_\sigma$ was also calculated by the change in the center of mass of cells per 1 USS. We ran the simulation until steps equivalent to 23 hours after a start of cell migration in the confinement release assay.

##### *4-2. Confinement release assay, Boundary conditions, and Initial conditions.*

In reference to a size of imaging windows, we set a pixel length of the simulation space as 2.5 μm × 2.5 μm, and the computer simulation was performed in a space with 1200 (horizontal) × 400 (vertical) pixels, corresponding to 3 mm × 1 mm. We placed a cluster of cells, arranged with 120 cells in the x-axis direction by 50 cells in the y-axis direction, in a simulation field with a reflecting boundary condition. There is a free space only along the left side of the cell cluster, but no free space along the top, bottom, or right side. We assumed that cells facing the free space were leader cells, so that leaders could be changeable throughout the simulation. At initiation, each cell area was set to the ideal area value, and the orientation of front-rear cell polarity in each cell was random except for the leader cells. Cell proliferation was not included in simulations because it seems to minimally affect the ERK activation wave propagation (Tlili et al., 2018).

##### *4-3. Leader cells and follower cells*

We defined the leader cells as cells facing a free space, corresponding to medium in the model, and follower cells as the others. The leader cells sense free spaces to migrate and communicate to the follower cells located behind the leader cells (Omelchenko et al., 2003; Yamaguchi et al., 2015). Thereby, we set the following two rules for the leader cells in polarity orientation and cell contraction. First, the cell polarity in the leader cells is persistently oriented towards the free space with a white Gaussian noise regardless of Eq. (4). Second, the leader cells keep generating polarized contractile force ($\omega = 1$) independent of cell extension-induced ERK activity ($\lambda_\sigma = 100$) in Eq. (2) and (3) throughout the simulations.

*4-4. Parameters*

The default parameter values are shown in the column "Values" in Table S1. In the rightmost column, the range of the respective parameters used in Fig. 8 is summarized (Table S1). Each parameter value was determined based on either measured data, observation, or a balance of other parameter values. Under an energy minimization framework, relative values of a parameter to values in other parameters should be significant if the system is not far from its equilibrium.

#### 5. Miscellaneous

Wave propagation of cell velocities and/or mechanical stress during collective cell migration have been modeled in some earlier studies, and most of them adopt a continuous approach (Alert and Trepat, 2019; Banerjee et al., 2015; Notbohm et al., 2016; Tlili et al., 2018; Yabunaka and Marcq, 2017). They are constructed with a relatively simple assumption and include just a few parameters, which makes the analysis easier. In contrast, we chose a cell-based approach to express the collective cell behavior and modeled a system on a cellular mechanochemical feedback at tissue scale. Despite being complex and limited to numerical simulations, our model can recapitulate multiple unidirectional ERK activation waves in collective cell migration. Moreover, it simulates conditions under which the waves are not generated, and those are consistent with our experimental results. We believe that this approach contributes to further understanding of unique systems within and between cells, as well as refining our theoretical modeling of such systems. Additional numerical investigations on multicellular dynamics and ERK activation waves were beyond the scope of this study. We will report on those elsewhere.

| Parameters/Variables | Symbols | Values [unit] | Range |
|---|---|---|---|
| Interfacial energy | $J_{cm}$,$J_{cc}$ | 3 | 2-4 |
| Magnitude of cell size constraint | $\lambda_A$ | 1, 0.02 | - |
| Ideal cell area | $A_0$ | 64 [pixels], 400 [$\mu m^2$] | - |
| Degree of polarity for contraction | $\omega$ | 1 | 0-1 |
| Polarity alignment to neighbors | $\mu$ | 1 | - |
| Timescale in ERK dynamics | $\eta_E$ | 5 [USS], 3 [min] | - |
| Sensitivity of ERK response to strain | $\alpha$ | 3 | - |
| Timescale in contraction dynamics | $\eta_c$ | 50 [USS], 30 [min] | - |
| Magnitude of maximum contraction | $\lambda$ | 10 | 0-10 |
| Threshold of ERK for contraction | $[ERK]^*$ | 0.5 | - |

**Table S1 Parameters used for simulations**
